# Intermolecular disulfide bond formation promotes Hsp42 higher-order assembly and shapes client selection in yeast

**DOI:** 10.64898/2026.07.13.738256

**Authors:** Long Duy Duong, Daniel Escobar-Osorio, Alexander B. Saltzman, Kevin A. Morano

## Abstract

Cellular redox homeostasis plays a critical role in regulating protein function, including chaperone activity, through reversible oxidation of cysteine and methionine residues. Previously, we found that budding yeast cells experiencing redox imbalance due to inactivated thioredoxin reductase (*trr1*Δ) activate the heat shock response and induce hyperaccumulation of the small heat shock protein/sequestrase Hsp42 with misfolded proteins. Building on that finding, this study identified cysteine 127 (C127) within Hsp42 as a redox-active residue that becomes oxidized in *trr1*Δ cells, upon treatment with the powerful oxidant hydrogen peroxide, or by exposure to the cysteine crosslinker divinyl sulfone (DVSF). In *trr1*Δ cells, C127 oxidation promoted intermolecular disulfide bond formation and contributed to Hsp42 homo-oligomerization. We show that stable oligomerization requires both the prion-like domain (PrLD) and C127 oxidation. While Hsp42–GFP formed prominent persistent foci in *trr1*Δ cells, replacement of C127 with non-thiol reactive serine decreased foci formation. Furthermore, the C127S mutation diminished Hsp42 oligomerization and sedimentability. Immunoprecipitation coupled with mass spectrometry analysis revealed that Hsp42 in *trr1*Δ cells preferentially associated with mitochondrial precursor proteins accumulated in the cytoplasm, as well as oxidation–reduction enzymes. The observed client selectivity was altered by the C127S mutation that diversified the spectrum of Hsp42-associated proteins. Collectively, these findings identify Cys127 as a redox-active switch that regulates Hsp42 assembly, foci formation, stability, and client specificity in response to oxidative stress.

## Introduction

Maintaining proteostasis, defined as a homeostatic balance of protein synthesis, folding, function, and degradation, is critical for life. It is particularly important under extrinsic or intrinsic proteotoxic stresses, including heat shock, oxidative stress, nutrient limitation, and aging, all of which can destabilize protein structure and promote protein misfolding and aggregation. Failed proteostasis can lead to the accumulation of toxic protein aggregates, impaired cellular functions, and ultimately cell death (1–4). To safeguard the proteome, cells employ an extensive protein quality control network relying largely on molecular chaperones that are highly conserved across all domains of life. Molecular chaperones include ATP-dependent “foldases”, such as Hsp70 and Hsp90, which facilitate protein folding and refolding; ATP-dependent disaggregases, such as Hsp104 and the Hsp110/Hsp70/Hsp40 machinery, both of which solubilize and extract aggregated proteins; and ATP-independent “holdases” and “sequestrases”, including the small heat shock proteins (sHSPs), that bind and/or sequester unfolded/misfolded proteins to maintain them in a folding-competent state (1, 3).

Among molecular chaperones, small heat shock proteins are considered a first line of defense against proteotoxic stress (5–7). They are characterized by a conserved α-crystallin domain (ACD), first identified in the eye lens protein α-crystallin (8, 9). The ACD is flanked by N-terminal (NTD) and C-terminal (CTD) regions that vary considerably in sequence and length among family members and contribute to their functional diversity (5). In response to proteotoxic stress, sHSPs assemble into dynamic oligomeric complexes through interactions among these domains. The ACD forms the structural core of sHSP oligomers, whereas the NTD and CTD regulate oligomer assembly, dynamics, and client interactions. sHSPs primarily recognize non-native proteins through exposed hydrophobic regions that become accessible upon protein unfolding or misfolding (5). Under conditions of elevated proteotoxic stress, cellular sHSP levels are rapidly increased and bind unfolded or misfolded proteins, thereby preventing their irreversible aggregation. In addition to acting as holdases, many sHSPs function as sequestrases that compartmentalize unfolded/misfolded proteins into specialized protein quality-control compartments (10–13). Sequestered substrates can subsequently be refolded by ATP-dependent chaperone systems, or targeted for degradation through the ubiquitin–proteasome system or autophagy (14, 15). Through these activities, sHSPs play critical roles in maintaining proteome integrity under adverse conditions (5, 14, 15).

While the budding yeast *Saccharomyces cerevisiae* contains only three α-crystallin domain-containing sHSPs (Btn2, Hsp26, and Hsp42), the sHSP family has expanded considerably in higher eukaryotes. Humans possess 10 sHSPs, while some plant species express as many as 40 (14–17). In budding yeast, Hsp26 and Hsp42 have been demonstrated to function as sequestrases that bind unfolded or misfolded proteins and direct them to transient cytoplasmic (CytoQs) or more persistent juxtanuclear quality control compartments (JUNQ) during proteotoxic stress (10, 14, 18, 19). In contrast, Btn2 promotes the sequestration of unfolded/misfolded proteins into the intranuclear quality-control compartment (INQ) (10, 14, 15). Beyond their roles in protein quality control during heat stress, Btn2 and Hsp42 have also been implicated in the oxidative stress response, where they contribute to hydrogen peroxide tolerance by compartmentalizing oxidized proteins into specialized deposition sites (20). Consistent with a role for Hsp42 in oxidative stress adaptation, we recently found that disruption of the cytosolic thioredoxin pathway, but not the glutathione pathway, leads to chronic accumulation specifically of Hsp42 into enlarged cytoplasmic foci, whereas Btn2 and Hs26 remain diffuse. This accumulation is accompanied by the sequestration of misfolded proteins, suggesting a dedicated role for Hsp42 in maintaining proteostasis under conditions of redox imbalance (19).

The thioredoxin pathway is the major cellular system that catalyzes the reduction of protein disulfide bonds, thereby modulating the folding, stability, activity, localization, and interactions of target proteins in budding yeast. As such, it serves as a key regulator of redox-dependent signaling and cellular processes. In contrast, the glutathione pathway functions predominantly as a cellular redox buffer and detoxification system that maintains general redox homeostasis (21–24). In *Saccharomyces cerevisiae*, the thioredoxin system comprises the thioredoxins Trx1, Trx2, and Trx3, and the NADPH-dependent thioredoxin reductases Trr1 and Trr2. Through sequential electron transfer from NADPH, thioredoxin reductases maintain thioredoxins in a reduced state, enabling them to reduce disulfide bonds in a broad range of target proteins (25). Trx1, Trx2, and Trr1 function predominantly in the cytosol, whereas Trx3 and Trr2 form the mitochondrial thioredoxin system that preserves mitochondrial redox homeostasis and plays an important role in respiratory growth when mitochondrial reactive oxygen species (ROS) production is elevated (26). In addition to promoting the accumulation of Hsp42 with misfolded proteins, disruption of the cytosolic thioredoxin system through deletion of *TRR1*, *TRX1*/*TRX2*, or all three genes results in a wide range of cellular defects including constitutive activation of Yap1-dependent antioxidant genes, elevated basal activation of the unfolded protein response (UPR), alteration in TORC1 signaling and autophagy, decreased tolerance to oxidative stress, and a shortened replicative lifespan (19, 27–31). The cytosolic thioredoxin system therefore plays a critical role in maintaining redox homeostasis and proteostasis.

Under oxidative stress, many proteins undergo reversible oxidative modifications, particularly at cysteine thiol groups and methionine thioether sulfur groups (32). These modifications can function as redox switches that alter protein structure and activity, thereby facilitating cellular adaptation to oxidative stress (4, 32, 33). Although Hsp42 forms stable and pronounced protein foci in redox-challenged *trr1*Δ cells, the mechanism by which Hsp42 may be regulated under oxidative stress remains unclear. In this study, we investigated the redox regulation of Hsp42 in cells lacking thioredoxin reductase. Hsp42 contains a single cysteine residue, Cys127, which we found to be redox-active and capable of forming intermolecular disulfide bonds, resulting in the formation of trimer-like homo-oligomeric species in *trr1*Δ cells. Formation of these oligomers requires both Cys127 and the prion-like domain (PrLD). Mutation of Cys127 to serine decreases Hsp42 oligomerization, foci formation, and sedimentability in *trr1*Δ cells. Furthermore, immunoprecipitation–mass spectrometry analysis revealed that Hsp42 preferentially associates with mislocalized mitochondrial and redox-related proteins in *trr1*Δ cells, whereas this client specificity is diminished by the C127S mutation. Together, these findings identify Cys127 as a redox-sensitive switch that regulates Hsp42 activity to promote the sequestration of mitochondrial and redox-related proteins during oxidative stress.

## Results

### Cysteine 127 of Hsp42 is redox sensitive and can form a disulfide bond in redox-challenged *trr1*Δ cells

Previously, we discovered that redox imbalance caused by disruption of the cytosolic thioredoxin pathway, but not the glutathione pathway, leads to accumulation of the small heat shock protein Hsp42 with misfolded proteins in an enlarged JUNQ compartment that fails to resolve over time (19). To further investigate how cellular redox stress regulates Hsp42 sequestration activity in *trr1*Δ cells, we focused on cysteine residues, as cysteine can exist in a redox-active thiolate anion form, and often functions as a regulatory switch under oxidative stress conditions (32). Interestingly, among the three small heat shock proteins, Hsp42 contains a single cysteine residue at position 127, located within its intrinsically disordered domain (IDD), whereas cysteine is absent in Btn2 and Hsp26 (Fig. 1A). We initially performed non-reducing SDS-PAGE analysis of wild-type (WT) and *trr1*Δ cells expressing all three sHSPs fused to GFP at their C-termini. Because the expression level of Btn2 is low in both WT and *trr1*Δ cells (19), we increased its expression using a multicopy plasmid to facilitate analysis. As shown in Fig. 1B, in WT cells, no additional species were detected for any of the three proteins beyond their monomeric forms. However, when expressed in *trr1*Δ cells and analyzed under non-reducing conditions, we detected a high-molecular-weight multimer of Hsp42, suggesting that Hsp42 can exist in a higher order, thiol-sensitive form in *trr1*Δ cells. Based on these observations, we hypothesized that the multimeric band may involve a disulfide bond at cysteine 127. To test this, we first sought to determine whether cysteine 127 is redox-active. We generated FLAG-tagged Hsp42 or Hsp42-C127S proteins that were immunoprecipitated using anti-FLAG M2 magnetic beads and treated with 10 kDa mPEG-MAL (methoxy polyethylene glycol maleimide) to assess cysteine reactivity. As shown in Fig. 1C, WT Hsp42 displayed a band shift following mPEG-MAL treatment, whereas the Hsp42-C127S mutant, which lacks a cysteine thiol group, did not. These results confirm that cysteine 127 is redox active. Consistently, mutation of cysteine 127 to serine, or treatment with the reducing agent β-mercaptoethanol, abolished Hsp42 multimer formation, indicating that Hsp42 forms intermolecular disulfide bonds in redox-challenged *trr1*Δ cells. (Fig. 1D). Together, these data suggest that redox perturbation caused by disruption of the cytosolic thioredoxin pathway promotes oxidation of Hsp42 at cysteine 127, leading to intermolecular disulfide bond formation.

**Figure 1.**
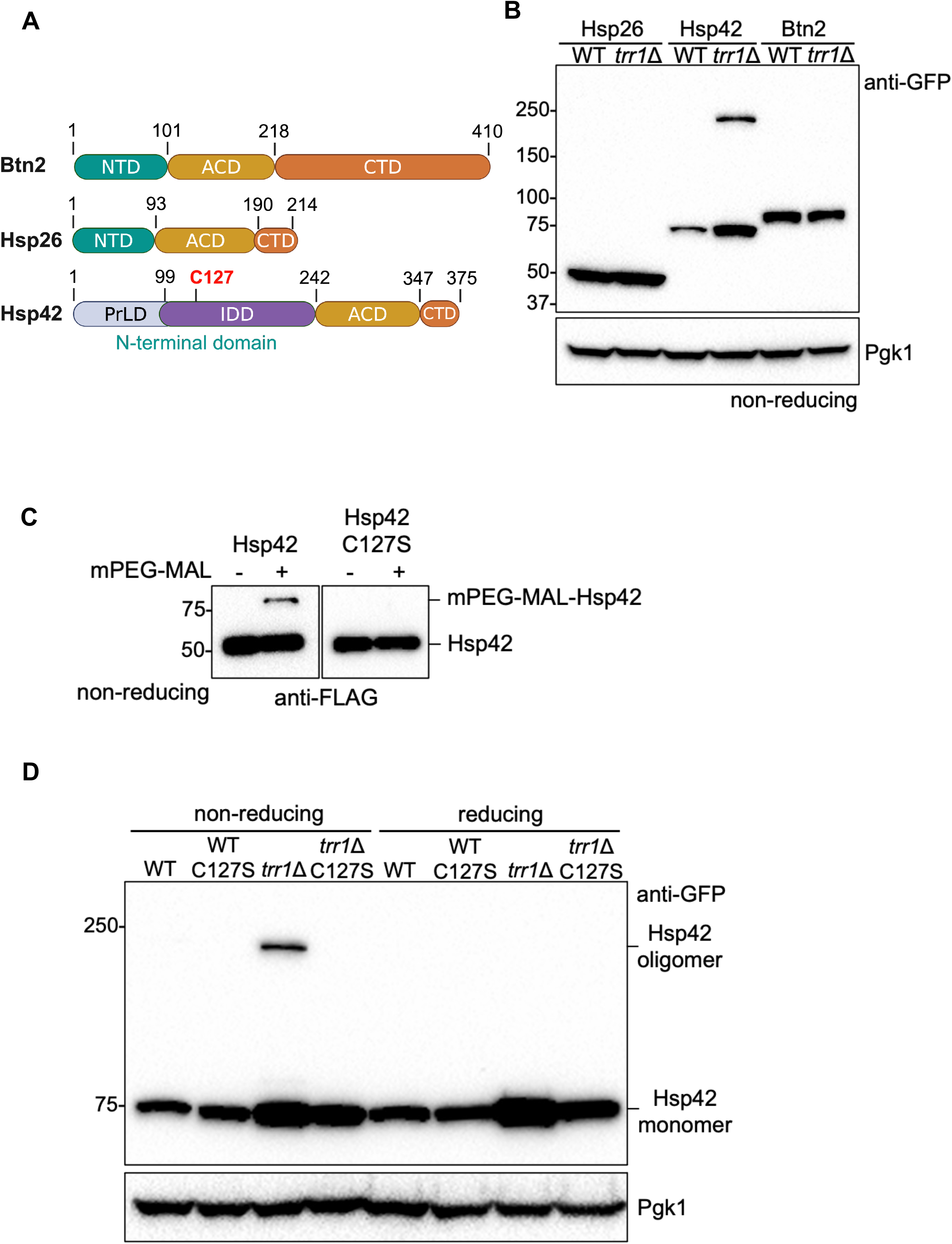
Hsp42 Cys127 is redox-active and forms disulfide bonds in *trr1*Δ cells. **(A)** Domain structures of the yeast small heat shock proteins Btn2, Hsp26, and Hsp42. **(B)** Hsp42-GFP from *trr1*Δ cells forms oligomeric species on non-reducing SDS-PAGE. WT and *trr1*Δ cells overexpressing Btn2-GFP were grown in SC-URA medium, whereas WT and *trr1*Δ cells expressing Hsp26-GFP or Hsp42-GFP were grown in YPD medium, until mid-log phase. Total protein extracts were prepared using the glass-bead lysis method and analyzed by non-reducing SDS-PAGE followed by Western blotting. **(C)** Hsp42 Cys127 is redox-active. Hsp42-FLAG and Hsp42-C127S-FLAG were immunoprecipitated from mid-log phase WT cells using anti-FLAG M2 affinity beads. The isolated proteins were incubated with 10-kDa mPEG-maleimide (mPEG-MAL) at room temperature for 30 min and subsequently analyzed by non-reducing SDS-PAGE followed by Western blotting. **(D)** Substitution of Cys127 with serine or treatment with a reducing agent abolishes Hsp42 oligomer formation. WT and *trr1*Δ cells expressing either Hsp42-GFP or Hsp42-C127S-GFP were grown to mid-log phase in YPD medium and then lysed using the glass-bead method. Cell lysates were subjected to either reducing SDS-PAGE containing 5% β-mercaptoethanol (β-ME) or non-reducing SDS-PAGE, followed by Western blot analysis.

### Thiol-modifying reagents modulate Hsp42 multimerization in WT cells

Since redox-induced Hsp42 multimerization occurs in redox-stressed *trr1*Δ cells, we next asked whether Hsp42 oligomerization also occurs under normal physiological conditions in the absence of elevated intracellular redox stress. Because disulfide bond formation is a dynamic process involving continuous oxidation and reduction of cysteine residues, we sought to determine whether Hsp42 disulfide linkages can also form in WT cells. To capture such transient oxidation events, we employed divinyl sulfone (DVSF), a thiol-reactive crosslinker that covalently links two closely positioned (8-10Å) thiols, thereby stabilizing dynamic or transient protein-protein interactions (34–36). To test this, WT cell lysates containing Hsp42-GFP were treated with DVSF. As shown in Fig. 2A, DVSF treatment induced Hsp42 oligomerization *in vitro*, as detected by non-reducing SDS-PAGE. Furthermore, oligomer formation increased in a dose-dependent manner with increasing DVSF concentrations. At 0.5 mM DVSF, nearly all detectable monomeric Hsp42-GFP was converted into oligomeric species. Consistent with these findings, treatment of live WT cells with DVSF also resulted in the accumulation of the same high-molecular-weight Hsp42 species (Fig. 2B), which was disrupted by C127S mutation (Fig. 3D), indicating that oxidation of Hsp42 at C127 occurs in WT cells under normal growth conditions. Together, these results suggest that Hsp42 oxidation at C127 is a dynamic process, and that DVSF traps Hsp42 in an oxidized state, thereby stabilizing oligomer formation even in the absence of redox stress.

**Figure 2.**
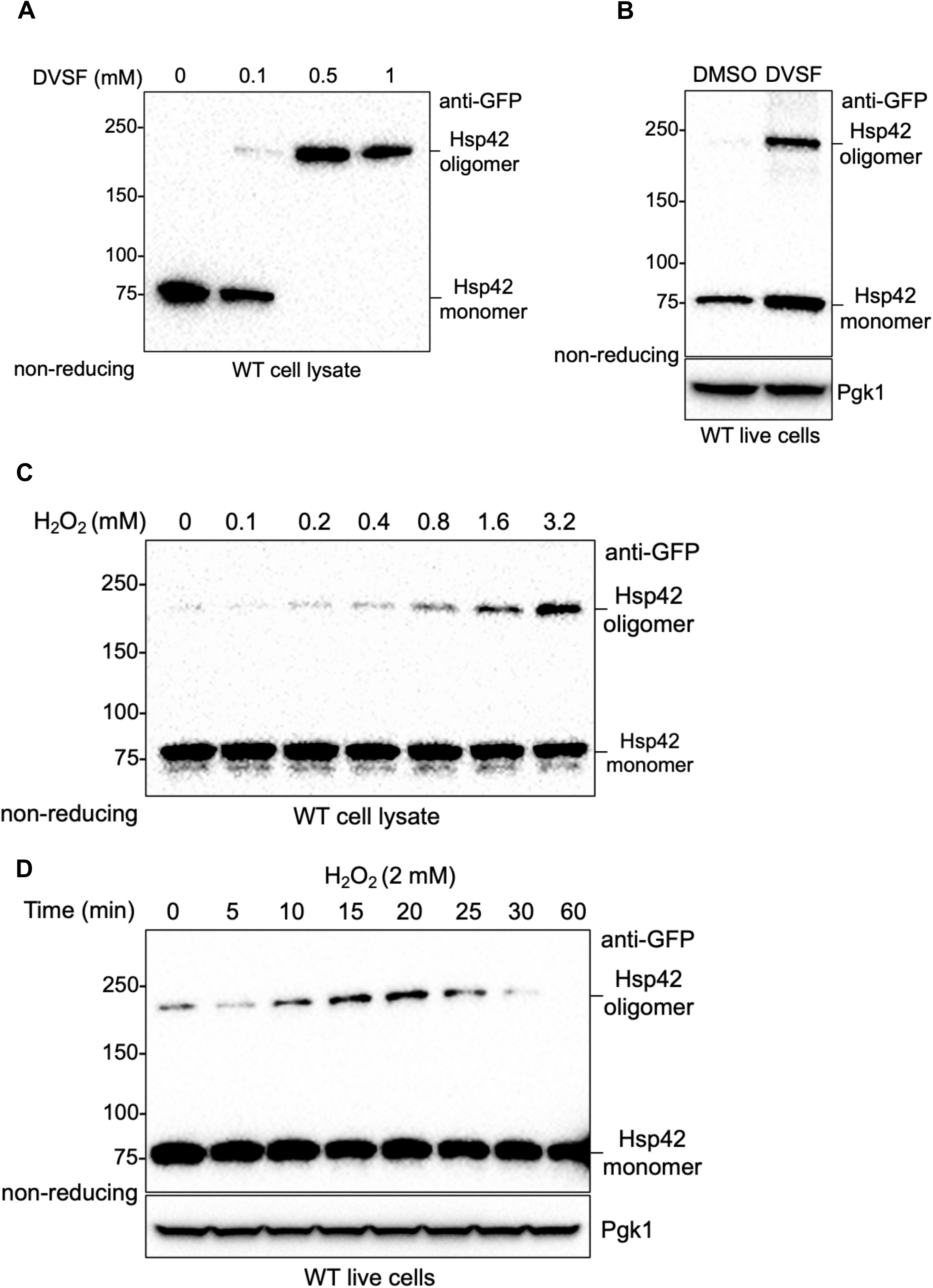
Thiol-modifying reagents promote Hsp42 multimerization in WT cells. (A) The thiol cross-linking agent DVSF induces Hsp42 oligomerization *in vitro*. Cell lysates prepared from mid-log phase WT cells expressing Hsp42-GFP by glass-bead lysis were treated with DVSF at the indicated concentrations (0.1, 0.5, or 1 mM) for 1 h. Samples were then analyzed by non-reducing SDS-PAGE followed by Western blotting. **(B)** DVSF promotes Hsp42 oligomerization *in vivo*. Mid-log phase WT cells expressing Hsp42-GFP were treated with 0.2 mM DVSF for 1 h before harvesting. Proteins extracted by glass-bead lysis were analyzed by non-reducing SDS-PAGE followed by Western blotting. **(C)** H_2_O_2_ induces Hsp42 oligomerization *in vitro*. Cell lysates prepared from mid-log phase WT cells expressing Hsp42-GFP by glass-bead lysis were treated with H_2_O_2_ at the indicated concentrations for 30 min at room temperature. Samples were then subjected to non-reducing SDS-PAGE followed by Western blotting. **(D)** H_2_O_2_ promotes Hsp42 oligomerization *in vivo*. Mid-log phase WT cells expressing Hsp42-GFP were treated with 2 mM H_2_O_2_ for the indicated times before harvesting. Protein extracts were analyzed by non-reducing SDS-PAGE followed by Western blotting.

**Figure 3.**
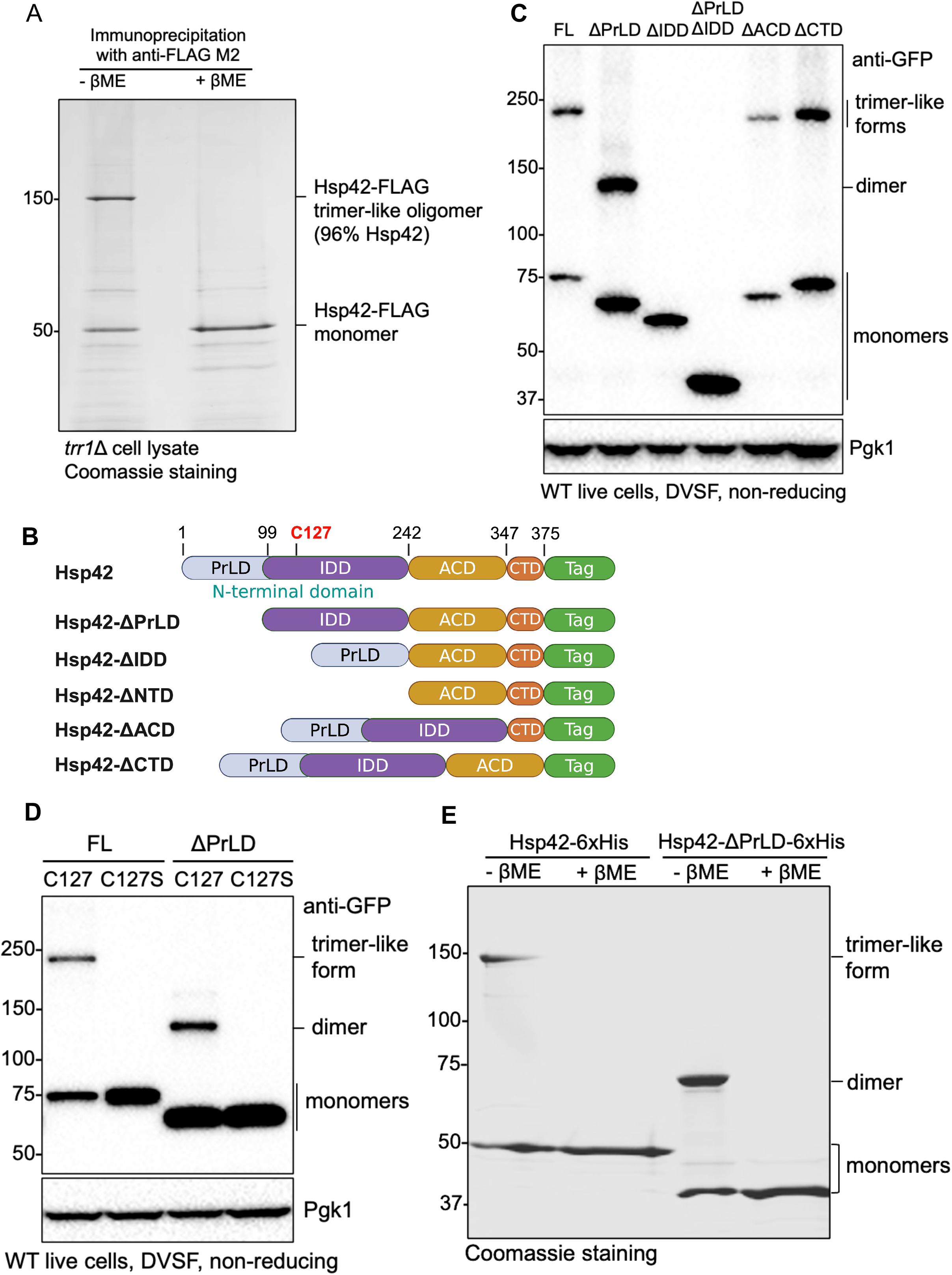
The prion-like domain and C127 are required for Hsp42 trimer-like homo-oligomer formation. **(A)** Hsp42 constitutes the majority (96%) of the trimer-like oligomeric species. Hsp42-FLAG was immunoprecipitated from *trr1*Δ cells and analyzed by SDS-PAGE followed by Coomassie staining. The band corresponding to the trimer-like oligomer was excised and subjected to mass spectrometry (MS) analysis. **(B)** Domain-deletion variants of Hsp42. **(C)** Effects of domain deletions on trimer-like oligomer formation of Hsp42. WT cells harboring plasmids expressing Hsp42-GFP(C48S,C70S) domain-deletion variants were grown in SC-Ura medium to mid-log phase and treated with 0.2 mM DVSF for 1 h prior to harvesting. Cell lysates were prepared using the glass-bead method and analyzed by non-reducing SDS-PAGE followed by Western blotting. **(D)** Cys127 is required for dimer formation of Hsp42-ΔPrLD. WT cells harboring plasmids expressing Hsp42-GFP(C48S,C70S), Hsp42-ΔPrLD-GFP(C48S,C70S), or their corresponding C127S variants were grown in SC-Ura medium to mid-log phase and treated with 0.2 mM DVSF for 1 h before harvesting. Cell lysates were prepared by glass-bead lysis and analyzed by non-reducing SDS-PAGE followed by Western blotting. **(E)** Hsp42-6xHis and Hsp42-ΔPrLD-6xHis expressed in *E. coli* exhibit similar oligomerization patterns on non-reducing SDS-PAGE. Recombinant Hsp42-6xHis and Hsp42-ΔPrLD-6xHis were purified from *E. coli* using HisPur™ cobalt resin and analyzed by SDS-PAGE in the presence or absence of reducing agent, followed by Coomassie staining.

The oxidative stress caused by *TRR1* deletion represents an endogenous source of redox stress. Previous studies have shown that treatment of WT cells with 0.8 mM hydrogen peroxide (H_2_O_2_) increases the formation of Hsp42-GFP foci (20). We therefore asked whether an exogenous oxidative challenge would induce the same Hsp42 multimerization pattern observed in *trr1*Δ cells. To determine whether H_2_O_2_ can directly oxidize Hsp42, we first treated WT cell extracts with increasing concentrations of H_2_O_2_. As shown in Fig. 2C, H_2_O_2_ treatment induced Hsp42 disulfide bond formation in a concentration-dependent manner, demonstrating that Hsp42 can be directly oxidized *in vitro* to form disulfide-linked oligomers. Next, we examined Hsp42 oligomerization *in vivo* by treating WT cells with H_2_O_2_ and monitoring oligomer formation over a 60-minute time course. As shown in Fig. 2D, Hsp42 oligomerization increased rapidly, reaching a maximum after 20 min of treatment. Thereafter, the abundance of oligomeric species gradually declined and was no longer detectable after 60 min. This transient response suggests that cellular antioxidant systems may be activated following H_2_O_2_ exposure, promoting the reduction of Hsp42 disulfide bonds. The subsequent decline in Hsp42 oligomers could reflect one or more processes, including H_2_O_2_ detoxification, oligomer dissociation upon reduction, or degradation of oxidized Hsp42 by cellular protein quality control pathways. Taken together, these results demonstrate that both endogenous and exogenous oxidative stress can induce Hsp42 oligomerization and that thiol-oxidizing agents promote the formation of disulfide-linked Hsp42 oligomers both *in vitro* and *in vivo*.

### The Prion-Like Domain (PrLD) influences Hsp42 multimer assembly

A sub-population of Hsp42-GFP in *trr1*Δ cells clearly exists as a higher-molecular-weight oligomer, as revealed using non-reducing SDS-PAGE. Given that monomeric Hsp42-GFP migrates at approximately 75 kDa, whereas the oligomeric species migrates at ∼225 kDa (Fig. 1B), the complex is unlikely to be a dimer and is instead consistent with a trimeric oligomer. A similar migration pattern was observed when the GFP tag was replaced with a FLAG tag (Fig. 3A), indicating that the oligomeric species is not an artifact of the GFP fusion. Because Hsp42 contains only a single cysteine residue, it remained possible that Hsp42 formed a disulfide bond with another cysteine-containing protein, resulting in a hetero-oligomeric complex. To distinguish between homo- and hetero-oligomer formation, we immunoprecipitated Hsp42-FLAG, the proteins were separated by non-reducing SDS-PAGE, and the oligomeric band was excised for mass spectrometry analysis. Hsp42 accounted for 96% of the protein identified within the oligomeric band, indicating that the complex nearly exclusively consists of Hsp42 and therefore represents a homo-oligomer. We refer to this species as a “trimer-like” form because its migration is consistent with that of a trimer, although it could alternatively represent a dimer with anomalous electrophoretic mobility under non-reducing conditions.

An interesting feature of Hsp42 is its N-terminal prion-like domain (PrLD), which has been predicted using a sequence-analysis approach and plays a critical role in substrate recognition and sequestration into cellular compartments during heat stress (37, 38). We therefore hypothesized that the PrLD contributes to formation of the trimer-like species. To test this idea, we generated a series of Hsp42 domain-deletion mutants (see Fig. 3B). Because GFP contains two cysteine residues (C48 and C70), which could potentially participate in unwanted disulfide bond formation, both residues were mutated to serine. Although this cysteine-free GFP variant lost fluorescence, it remained stable and readily detectable by immunoblotting. WT cells expressing the domain-deletion Hsp42-GFP(C48S,C70S) plasmids were treated with DVSF to induce formation of the trimer-like species. As shown in Fig. 3C, deletion of either the ACD or CTD did not affect formation of the trimer-like oligomers. In contrast, deletion of the IDD or the entire NTD (PrLD and IDD) completely abolished oligomer formation. Because C127, the residue responsible for disulfide bond formation, resides within the IDD, loss of the oligomer in these mutants was expected. Interestingly, deletion of the PrLD alone did not eliminate oligomerization. Instead, the mutant produced a different higher-molecular-weight oligomeric species that migrated at approximately the size expected for a dimer rather than the trimer-like form observed in full-length Hsp42. Similar to the trimer-like species, formation of this dimeric complex was abolished by the C127S mutation (Fig. 3D). These findings suggest that, in addition to C127 oxidation, the PrLD plays an important role in determining the oligomeric architecture of Hsp42.

To exclude the possibility that other post-translational modifications contribute to formation of the trimer-like species, we expressed and purified Hsp42-6xHis and Hsp42-ΔPrLD-6xHis from *E. coli*. Consistent with the results obtained in yeast, purified Hsp42-6xHis formed the trimer-like species, whereas Hsp42-ΔPrLD-6xHis predominantly formed a dimeric species (Fig. 3E). Importantly, both the trimer-like and dimeric forms were abolished by treatment with the reducing agent β-mercaptoethanol (β-ME), confirming that their formation depends on disulfide bond formation. These observations further confirm that the distinct oligomerization patterns are intrinsic properties of Hsp42 and do not require yeast-specific post-translational modifications. Taken together, these results demonstrate that C127-mediated disulfide bond formation drives Hsp42 oligomerization, whereas the PrLD plays a critical role in promoting assembly of the higher-order trimer-like species.

### The PrLD and cysteine 127 regulate Hsp42 foci formation in redox challenged *trr1*Δ cells

Previous studies have shown that the PrLD of Hsp42 plays a critical role in formation of consolidated foci during heat shock, whereas IDD deletion results in multiple smaller foci (38). We therefore sought to determine the contribution of each domain to Hsp42-GFP foci formation in *trr1*Δ cells by generating a series of endogenous domain-deletion mutants as illustrated in Fig. 3B. First, we examined the steady-state protein levels of endogenously expressed Hsp42-GFP domain deletion variants in *trr1*Δ cells by immunoblotting. As shown in Fig. 4A, deletion of the PrLD, IDD, or the entire NTD had little effect on Hsp42-GFP protein abundance compared with full-length Hsp42-GFP. In contrast, deletion of either the ACD or CTD reduced protein stability, with ACD deletion producing the most pronounced decrease in protein levels.

**Figure 4.**
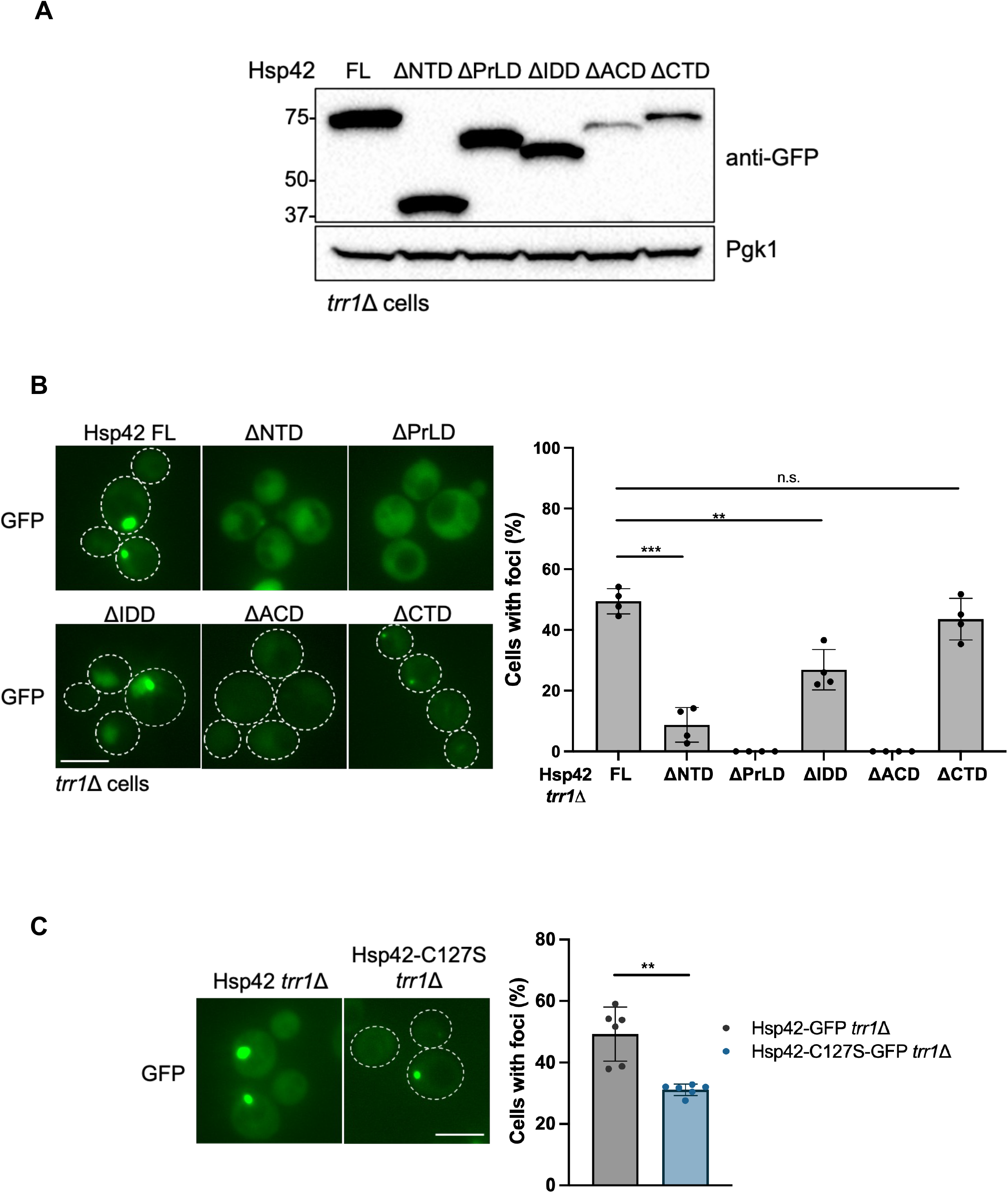
The PrLD is required for Hsp42-GFP foci formation, whereas Cys127 promotes foci formation in *trr1*Δ cells. **(A)** Expression levels of Hsp42-GFP domain-deletion variants in *trr1*Δ cells. The *trr1*Δ cells expressing Hsp42-GFP full-length (FT) and Hsp42-GFP domain-deletion variants from the endogenous locus were grown in YPD medium to mid-log phase and lysed using the glass-bead method. The protein extracts were analyzed by reducing SDS-PAGE followed by Western blotting. **(B)** The PrLD is critical for Hsp42-GFP foci formation in *trr1*Δ cells, and **(C)** the C127S mutation reduces Hsp42-GFP foci formation in *trr1*Δ cells. The *trr1*Δ cells expressing Hsp42-GFP, Hsp42-GFP domain-deletion variants, or Hsp42-C127S-GFP from the endogenous locus were grown in YPD medium to mid-log phase and examined for GFP fluorescence by microscopy. For quantification, at least four biological replicates were analyzed, with a minimum of three fields and 100 cells counted per replicate. The percentage of cells containing GFP foci was determined. Statistical significance between the indicated strains was assessed using an unpaired Student’s *t*-test (*p* < 0.05, *; *p* < 0.01, **; *p* < 0.001, ***; n.s., not significant). Scale bar, 5 μm.

Next, we examined the ability of these mutants to form intracellular foci in *trr1*Δ cells. As shown in Fig. 4B, deletion of the PrLD completely abolished Hsp42 foci formation, resulting in a diffuse GFP signal throughout the cytoplasm, indicating that the PrLD is essential for Hsp42 assembly into foci. Consistent with this observation, deletion of the entire NTD also produced a diffuse GFP localization pattern, with slightly stronger fluorescence in a region possibly corresponding to the nucleus, likely because this construct lacks the PrLD. Deletion of the ACD similarly eliminated foci formation; however, this effect is likely attributable to the severe drop in protein abundance observed by immunoblotting. In contrast, deletion of the CTD did not abolish foci formation. Although Hsp42-ΔCTD-GFP was expressed at lower levels than the full-length protein, it still formed distinct smaller puncta in cells. Moreover, the percentage of cells containing foci was comparable to that observed for full-length Hsp42-GFP, suggesting that the CTD is largely dispensable for foci formation. Interestingly, deletion of the IDD significantly reduced the proportion of cells containing Hsp42 foci compared with full-length Hsp42-GFP. In addition, Hsp42-ΔIDD-GFP exhibited a diffuse fluorescence signal that also seemed to be enriched within the nucleus. These findings suggest that the IDD contributes to the regulation of Hsp42 foci formation under conditions of redox stress.

Because C127 within the IDD undergoes oxidation and disulfide bond formation in *trr1*Δ cells, we next examined the effect of the C127S mutation on Hsp42-GFP foci formation. As shown in Fig. 4C, the C127S mutation significantly reduced the percentage of *trr1*Δ cells containing Hsp42-GFP foci, although it did not completely abolish foci formation. These results suggest that the PrLD serves as the primary determinant of Hsp42 assembly into foci, whereas C127 oxidation provides an additional regulatory mechanism that enhances foci formation under redox stress conditions. Taken together, these findings demonstrate that both the PrLD and C127 contribute to Hsp42 foci formation in redox-challenged *trr1*Δ cells, with the PrLD playing a dominant role and C127 oxidation acting as an auxiliary regulatory factor.

### The C127S mutation decreases Hsp42 oligomer size and sedimentability

Because C127 undergoes oxidation and disulfide bond formation in redox-challenged *trr1*Δ cells, resulting in the appearance of homo-trimer-like Hsp42 oligomers on non-reducing SDS-PAGE, we next sought to determine whether these oligomeric species could also be detected under native conditions. To address this question, we expressed Hsp42-6xHis and Hsp42-C127S-6xHis in *E. coli* and purified the protein in a soluble form (Supplementary Fig. S1). To ensure equivalent protein loading, purified protein concentrations were determined and analyzed by reducing SDS-PAGE followed by Coomassie staining (Supplementary Fig. S2) before being subjected to blue native PAGE (BN-PAGE) alongside native molecular-weight markers. As shown in Fig. 5A, Hsp42-6xHis migrated as multiple oligomeric species on BN-PAGE. However, neither dimeric nor trimeric species were readily detected, in contrast to the trimer-like band observed by non-reducing SDS-PAGE (Fig. 3E). These findings suggest that Hsp42 predominantly exists as large oligomeric assemblies in solution and that the apparent monomeric and trimer-like species observed on non-reducing SDS-PAGE likely arise from dissociation of these higher-order complexes in the presence of SDS. Indeed, the presence of multiple high-molecular-weight species on BN-PAGE indicates that Hsp42 is highly polydisperse and capable of assembling into large oligomeric structures, as previously observed (39, 40).

**Figure 5.**
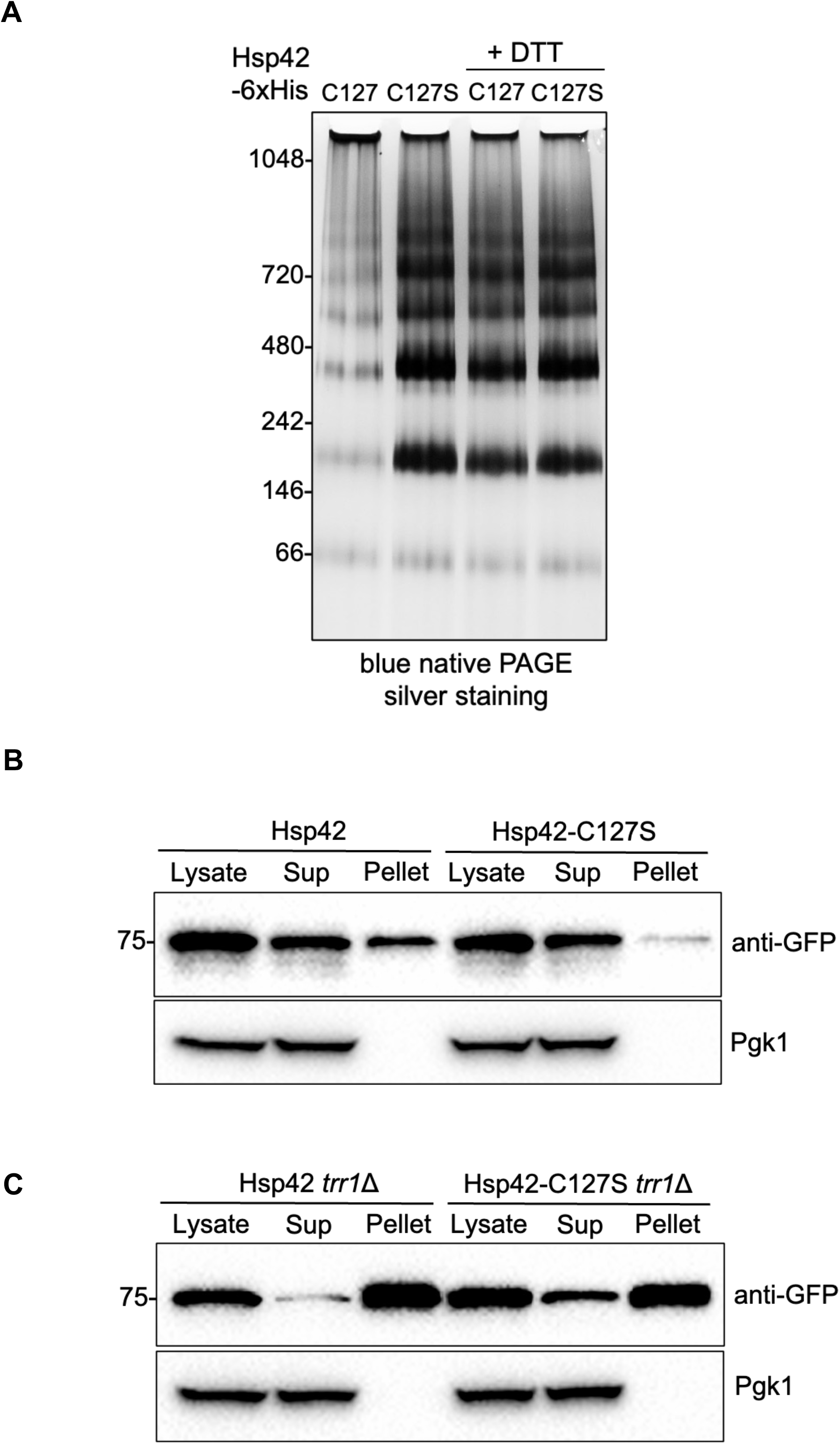
The C127S mutation decreases Hsp42 oligomer size and sedimentability. **(A)** The C127S mutation increases the abundance of lower-molecular-weight Hsp42 oligomers. Recombinant Hsp42-6xHis and Hsp42-C127S-6xHis proteins were purified from *E. coli*, and 3 μg of each protein was analyzed by blue native PAGE in the presence or absence of 10 mM DTT. Protein bands were visualized by silver staining. Fractionation analysis of WT cells **(B)** and *trr1*Δ cells **(C)** expressing Hsp42-GFP or Hsp42-C127S-GFP. The WT and *trr1*Δ cells expressing endogenous Hsp42-GFP or Hsp42-C127S-GFP were grown in YPD medium to mid-log phase, harvested, and fractionated as described in the Materials and Methods. Protein fractions were analyzed by reducing SDS-PAGE followed by Western blotting.

To determine whether C127-mediated disulfide bond formation contributes to this oligomeric organization, we also analyzed purified Hsp42C127S-6xHis by BN-PAGE. As shown in Fig. 5A, the C127S mutation increased the abundance of lower-molecular-weight oligomeric species relative to WT Hsp42-6xHis, suggesting that disruption of disulfide bond formation shifts the oligomeric equilibrium toward smaller assemblies. This observation is consistent with a model in which C127-dependent disulfide bonds act as molecular linkers that promote the association of smaller oligomers into larger complexes. To further test the role of C127-C127 intermolecular disulfide bonds, purified Hsp42-6xHis was treated with the reducing agent DTT prior to BN-PAGE analysis. Similar to the C127S mutation, DTT treatment increased the abundance of lower-molecular-weight oligomers (Fig. 5A), supporting the conclusion that disulfide bonds contribute to the stabilization of higher-order Hsp42 assemblies. In contrast, DTT treatment had little effect on the oligomeric profile of Hsp42-C127S-6xHis (Fig. 5A), confirming that the observed effect of DTT is mediated through C127. Taken together, these results demonstrate that Hsp42 exists as a heterogeneous population of oligomeric assemblies under native conditions, and that intermolecular disulfide bond formation at C127 promotes the assembly and stabilization of higher-order Hsp42 oligomers by facilitating interactions between smaller oligomeric species.

Based on these findings, we hypothesized that C127-mediated disulfide bond formation promotes the assembly of very large (> 1 MDa) Hsp42 oligomers at the expense of smaller oligomeric species, thereby increasing Hsp42 sedimentability. To test this hypothesis, we performed biochemical fractionation of WT and *trr1*Δ cells expressing either Hsp42-GFP or Hsp42-C127S-GFP and examined the distribution of Hsp42 between the supernatant and particulate fractions following centrifugation of cell lysates. In WT cells (Fig. 5B), Hsp42-GFP was detected in both the supernatant and pellet fractions, whereas the majority of Hsp42-C127S-GFP was recovered in the supernatant fraction. In *trr1*Δ cells (Fig. 5C), the majority of wild-type Hsp42-GFP was recovered in the pellet fraction, while a substantial proportion of Hsp42-C127S-GFP remained in the supernatant fraction, consistent with decreased protein sedimentability as governed by particle size. The redistribution of Hsp42-GFP from the supernatant to the pellet fraction in *trr1*Δ cells (Fig. 5B,C) was consistent with the increased Hsp42-GFP foci formation observed relative to WT cells as shown in our previous report (19). In contrast, the C127S mutation increased the proportion of Hsp42-GFP recovered in the *trr1*Δ supernatant fraction (Fig. 5C) and concomitantly decreased Hsp42 foci formation (Fig. 4C). These results support the idea that C127-mediated disulfide bond formation promotes the assembly of higher-order oligomeric structures in *trr1*Δ cells.

### Hsp42 client proteins are enriched for mitochondrial and redox-related proteins in ***trr1*Δ cells**

Because Hsp42 forms persistent protein foci that are associated with misfolded proteins in redox-challenged *trr1*Δ cells, we sought to identify Hsp42 client proteins under these conditions using immunoprecipitation coupled with mass spectrometry (IP-MS). To perform the immunoprecipitation, we used GFP-Trap magnetic beads to isolate Hsp42-GFP, or GFP alone as a negative control, from WT and *trr1*Δ cells. The control GFP protein expressed endogenously from the *HSP42* promoter showed diffuse fluorescence signals in both WT and *trr1*Δ cells (Supplementary Fig. S3). As shown in Fig. 6A, Hsp42-GFP and GFP were efficiently recovered and subsequently subjected to mass spectrometric analysis. Proteins exhibiting at least an eightfold enrichment relative to the GFP control and a *p*-value < 0.05 were considered significant interactors. Using these criteria, we identified 87 Hsp42-associated proteins in WT cells (Fig. 6B, Supplementary Table S2) and 173 Hsp42-associated proteins in *trr1*Δ cells (Fig. 6C, Supplementary Table S4). Remarkably, only nine proteins were shared between the two datasets, indicating extensive remodeling of the Hsp42 interactome upon loss of *TRR1*. Thus, deletion of *TRR1* increased the number of Hsp42-associated proteins by more than twofold, with approximately 95% of the interactors in *trr1*Δ cells representing newly acquired client proteins (Fig. 6E).

**Figure 6.**
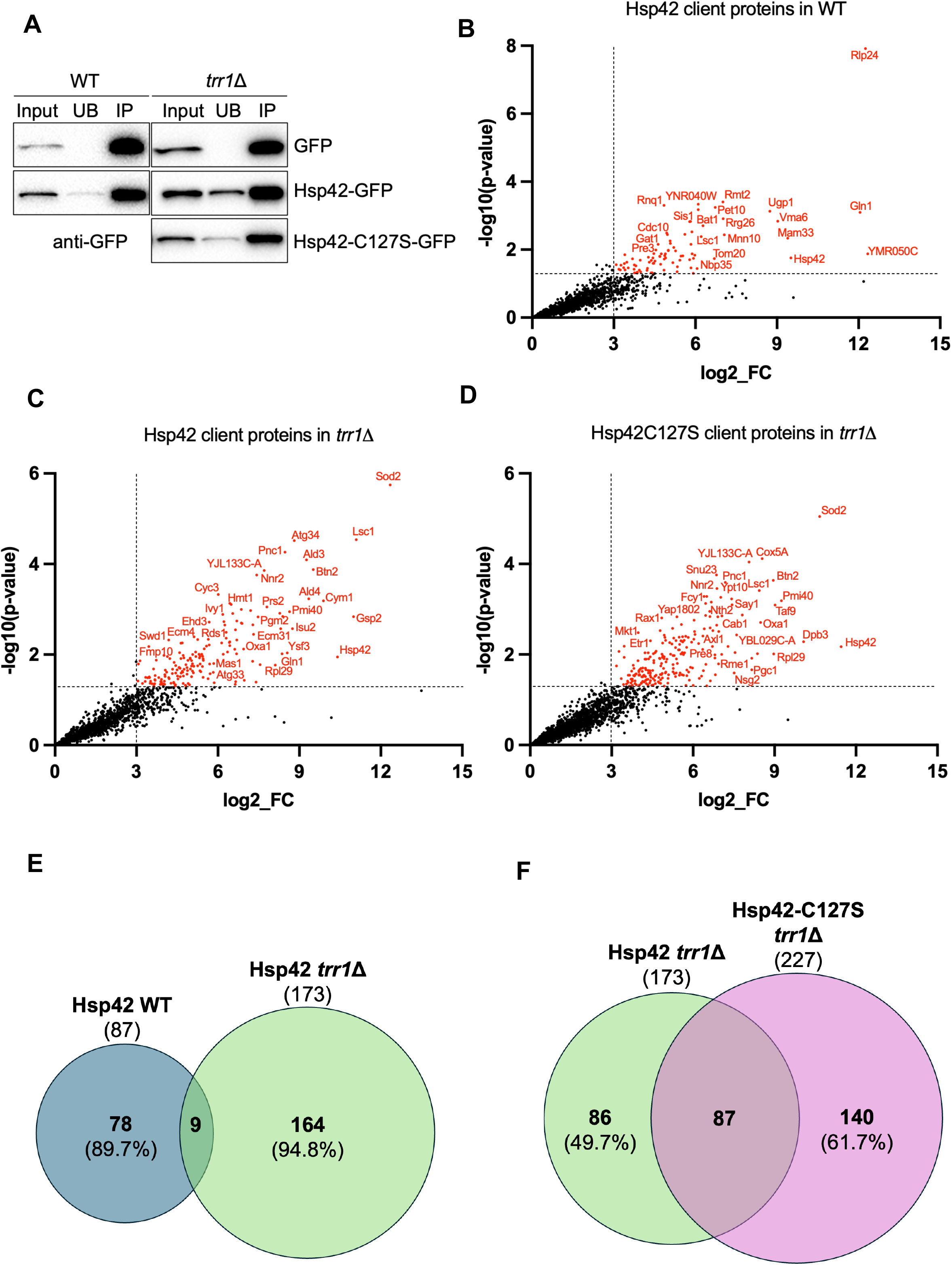
Client proteins of Hsp42-GFP in WT and *trr1*Δ cells and of Hsp42-C127S-GFP in *trr1*Δ cells. **(A)** Immunoprecipitation of GFP, Hsp42-GFP, and Hsp42-C127S-GFP using GFP-Trap. Cells were grown in YPD medium to mid-log phase, harvested, and subjected to immunoprecipitation as described in the Materials and Methods. Total cell lysate (Input), unbound (UB), and immunoprecipitated (IP) fractions were analyzed by reducing SDS-PAGE followed by Western blotting. **(B)** Client proteins associated with Hsp42-GFP in WT cells. **(C)** Client proteins associated with Hsp42-GFP in *trr1*Δ cells. **(D)** Client proteins associated with Hsp42-C127S-GFP in *trr1*Δ cells. For panels B–D, immunoprecipitated samples were analyzed by mass spectrometry to identify enriched proteins as described in the Materials and Methods with two biological replicates for each condition. **(E)** Number of shared client proteins associated with Hsp42-GFP in WT and *trr1*Δ cells. **(F)** Number of shared client proteins associated with Hsp42-GFP and Hsp42-C127S-GFP in *trr1*Δ cells.

To gain insight into the functional properties of these client proteins, we performed gene ontology (GO) enrichment analyses for cellular component, biological process, and molecular function. In WT cells, Hsp42-GFP-associated proteins were significantly enriched for components of the proteasome complex, including Pre3, Pre5, Pre8, Rpn3, and Rpn9, as well as enzymes involved in the tricarboxylic acid (TCA) cycle, such as Fum1, Idh1, Idh2, and Lsc1 (Fig. 7A, Supplementary Table S3). In contrast, Hsp42-GFP-associated proteins identified in *trr1*Δ cells were strongly enriched for mitochondrial proteins, such as Ald4, Coq6, Cta1, Mas1, Oxa1, Sam37, Sod2, and Tim50 which collectively accounted for approximately one-third of the associated proteins (59/173; Fig. 7B, Supplementary Table S5). Furthermore, 22 of the 173 associated proteins were linked to oxidoreductase activity, including Ald3, Ald4, Cta1, Ctt1, Oye2, Sod2, Zwf1, and Zta1. These findings suggest that deletion of *TRR1* substantially increases the burden of misfolded or aggregation-prone proteins involved in mitochondrial function and cellular redox homeostasis, thereby expanding the network of proteins associated with Hsp42.

**Figure 7.**
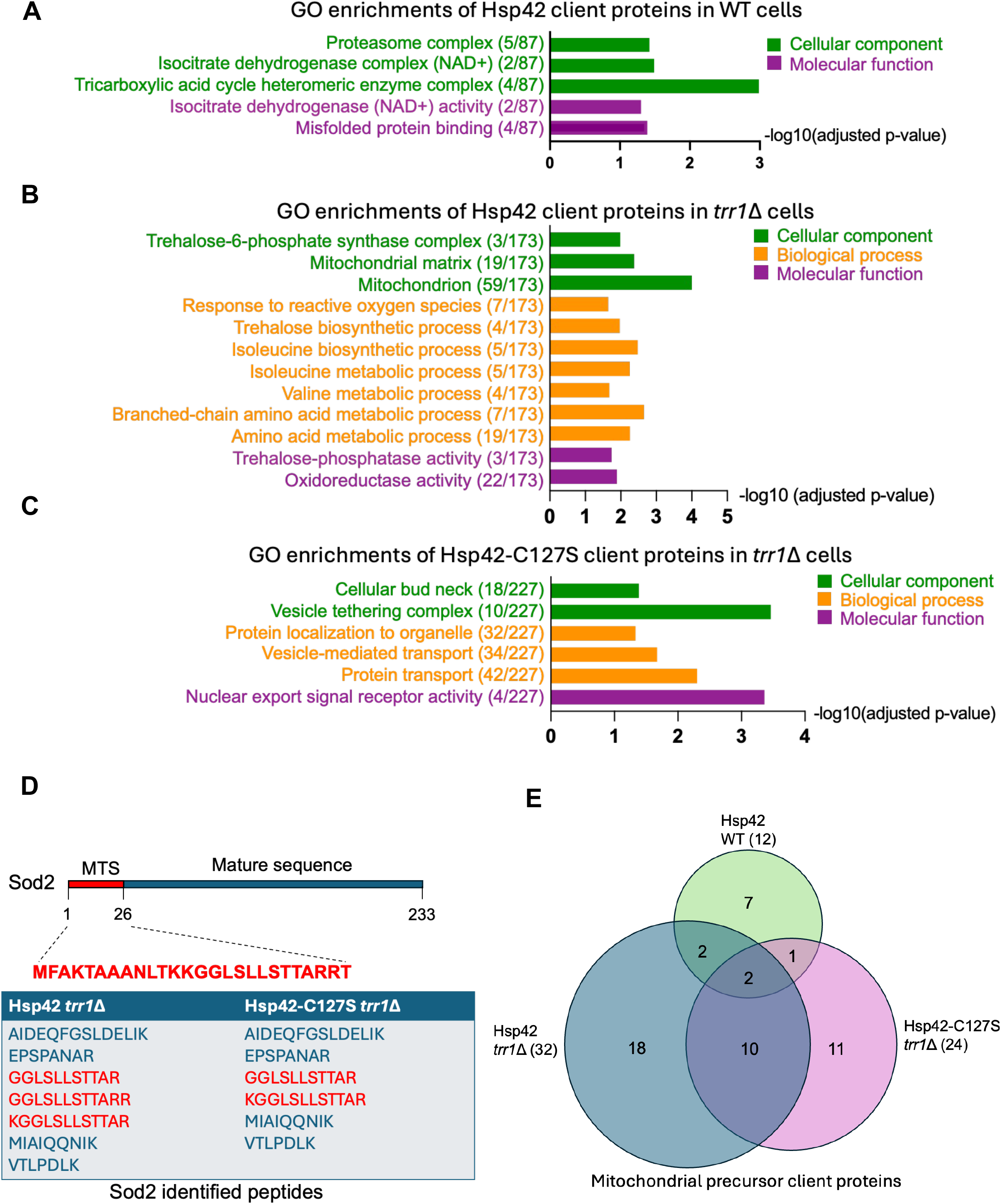
Gene ontology enrichment analysis of Hsp42-associated client proteins. **(A)** Gene ontology (GO) enrichment analysis of Hsp42-GFP-associated proteins in WT cells. **(B)** GO enrichment analysis of Hsp42-GFP-associated proteins in *trr1*Δ cells. **(C)** GO enrichment analysis of Hsp42-C127S-GFP-associated proteins in *trr1*Δ cells. GO enrichment analyses were performed using g:Profiler as described in the Materials and Methods. **(D)** Detection of Sod2-derived peptides in Hsp42-GFP and Hsp42-C127S-GFP immunoprecipitates from *trr1*Δ cells. **(E)** Number of mitochondrial precursor proteins associated with Hsp42-GFP in WT and *trr1*Δ cells and with Hsp42-C127S-GFP in *trr1*Δ cells.

### C127S mutation broadens client interactions and decreases selectivity in *trr1*Δ cells

Next, we investigated whether the decrease in Hsp42-GFP foci formation and large oligomer stability caused by the C127S mutation affect the interactome of Hsp42-GFP in redox challenged *trr1*Δ cells by IP-MS using Hsp42-C127S-GFP. As shown in Fig. 6A, Hsp42-C127S-GFP was also efficiently enriched using GFP-Trap magnetic beads. Using the same criteria applied to Hsp42-GFP, we identified 227 Hsp42-C127S-associated proteins (Fig. 6D, Supplementary Table S6), substantially more than the 173 client proteins identified for Hsp42-GFP, in *trr1*Δ cells. Comparison of the two datasets revealed that only 87 proteins were shared between WT Hsp42-GFP and Hsp42-C127S-GFP (Fig. 6F). Thus, approximately 50% of the WT Hsp42-associated proteins were retained in the C127S mutant, whereas the mutant acquired 140 additional interactors, representing approximately 62% of the total Hsp42-C127S-associated protein set. These findings indicate that the C127S mutation markedly expands the spectrum of proteins associated with Hsp42.

Enrichment analysis of the Hsp42-C127S-associated protein set revealed significant over-representation of proteins associated with the bud neck and protein transport pathways (Fig. 7C, Supplementary Table S7). In contrast, mitochondrial proteins and oxidoreductase enzymes were not significantly enriched, despite comprising 58 of 227 and 19 of 227 associated proteins, respectively (Supplementary Table S7). The loss of statistical significance for these categories likely reflects the broader and more diverse client repertoire acquired by the C127S mutant, which dilutes the relative contribution of mitochondrial and redox-related proteins. In summary, these results suggest that C127-mediated disulfide bond formation contributes to the specificity of Hsp42-client interactions under redox stress condition, potentially by influencing size and stability of the Hsp42 large multimeric forms. Disruption of this regulatory mechanism by the C127S mutation broadens the spectrum of Hsp42-associated proteins while lowering its preferential association with mitochondrial and redox-related client proteins.

### Precursor mitochondrial proteins associate with Hsp42 in *trr1*Δ cells

Among the proteins identified by IP-MS of Hsp42-GFP and Hsp42-C127S-GFP in *trr1*Δ cells, one protein stood out because of its high enrichment and extremely low *p*-value: Sod2, the major mitochondrial matrix antioxidant enzyme. Although Sod2 functions in mitochondria, it is encoded by a nuclear gene and synthesized in the cytosol as a precursor protein containing a 26-amino-acid N-terminal mitochondrial targeting sequence (MTS), which is cleaved following import into the mitochondrial matrix (41). This observation raised an intriguing question: why does Hsp42, a cytosolic chaperone, associate with a mitochondrial protein? To address this, we examined the Sod2-derived peptides identified by mass spectrometry in *trr1*Δ cells expressing either Hsp42-GFP or Hsp42-C127S-GFP. Surprisingly, peptides corresponding to the mitochondrial targeting sequence (MTS) region were detected and enriched in both datasets (Fig. 7D), indicating that Hsp42 associates with the precursor form of Sod2 rather than the mature mitochondrial protein.

We next extended this analysis to all mitochondrial proteins associated with Hsp42 in all IP-MS samples. Specifically, we searched for mitochondrial proteins whose MTS regions have been experimentally verified or predicted using MTSviewer (42). Given the cytoplasmic localization of Hsp42, its associated mitochondrial proteins are likely to represent precursor proteins before their import into mitochondria. As shown in Fig. 7E and Supplementary Table S8, only 12 mitochondrial precursor proteins were associated with Hsp42-GFP in WT cells. This number increased markedly to 32 in *trr1*Δ cells, consistent with elevated mitochondrial import stress under redox stress. Notably, the number of precursor proteins associated with Hsp42 was reduced to 24 in *trr1*Δ cells expressing Hsp42-C127S-GFP. Among the 51 mitochondrial precursor proteins identified across all tested conditions, MTS-containing peptides of 16 proteins (including Sod2) were recovered from IP-MS, with most of these peptides detected in *trr1*Δ cells (Supplementary Data 1). Taken together, these findings suggest that loss of *TRR1* impairs mitochondrial protein import, leading to the accumulation of mitochondrial precursor proteins in the cytosol, followed by sequestration by Hsp42. Furthermore, the decrease observed in the C127S mutant suggests that C127-mediated disulfide bond formation also contributes to the recognition by Hsp42 of mitochondrial precursor proteins under oxidative stress.

## Discussion

Oxidative stress, whether arising from external insults such as H₂O₂ exposure or internal perturbations such as disruption of redox pathways, compromises protein homeostasis by oxidizing proteins and impairing their functions (4). Several ATP-dependent molecular chaperones such as Hsp70s and Hsp90s have been shown to function as redox switches, whereby oxidation converts an ATP-dependent foldase into an ATP-independent holdase (4, 32, 33). In contrast, redox regulation of sHSPs has received considerably less attention although they are numerous in higher eukaryotes. Among the three yeast sHSPs, Hsp42 is unique in containing a single cysteine residue, Cys127. Our data demonstrate that this residue is redox-active, as evidenced by its reactivity toward thiol-modifying reagents including mPEG-MAL, DVSF, and H_2_O_2_. Using non-reducing SDS-PAGE, we observed the formation of disulfide-linked Hsp42 species in *trr1*Δ cells. These species were abolished either by mutation of Cys127 to serine or by treatment of lysates with reducing agents, indicating that oxidation of Cys127 promotes disulfide bond formation. Together, these findings establish Hsp42 as a redox-sensitive chaperone that undergoes cysteine oxidation under conditions of redox stress. Interestingly, oxidized C127 could also be detected in wild-type cells grown under non-stress conditions when cells were treated with the thiol cross-linker DVSF. Because DVSF traps proximal thiols and stabilizes oxidized thiol states (34–36), these observations suggest that Hsp42 may undergo transient oxidation even during normal growth, although such species are rapidly reduced under physiological conditions.

Oxidation of Hsp42 was not restricted to internally generated redox stress but was also observed following exposure to the exogenous oxidant H_2_O_2_. Both *in vitro* and *in vivo* experiments demonstrated Hsp42 oxidized in WT cells upon H_2_O_2_ treatment. *In vivo*, Hsp42 oxidation reached a maximum approximately 20 min after oxidant exposure but subsequently declined and became undetectable by 60 min. This transient oxidation profile suggests that cellular antioxidant programs regulated by Yap1 and Skn7 transcription factors (43) may be activated following oxidative challenge and efficiently restore the reduced state of Hsp42. Early oxidation of Hsp42 may play an important role in maintaining proteostasis by providing a rapid protective mechanism that preserves protein homeostasis while antioxidant defense genes are being activated. Collectively, our findings identify Hsp42 as a redox-sensitive small heat shock protein whose cysteine residue undergoes reversible oxidation in response to oxidative stress. These results support a model in which Hsp42 functions not only as a sensor of protein misfolding but also as a direct responder to changes in cellular redox status. This mechanism may be evolutionarily conserved from yeast to mammals, as 9 of the 10 canonical small heat shock proteins (sHSPs) in both rats and humans contain at least one cysteine residue. Supporting this possibility, Cys137 in HSPB1, which undergoes oxidation, is required for cellular protection against oxidative injury in rat cardiomyocytes and contributes to the anti-apoptotic activity in oxidatively stressed HeLa cells (44, 45).

Under stress conditions such as heat shock, Hsp42 assembles into oligomeric complexes that bind unfolded or misfolded proteins. These oligomers exhibit considerable structural heterogeneity and can exist in a wide range of sizes and architectures. For example, cell fractionation coupled with size-exclusion chromatography identified multiple distinct Hsp42-containing complexes larger than 200 kDa (39). In another report, blue native PAGE (BN-PAGE) analysis of Hsp42 purified from yeast revealed Hsp42 oligomers ranging from approximately 600–700 kDa to more than 1100 kDa (40). Given the highly polydisperse nature of Hsp42 oligomers, an important question is how oxidation of Cys127 influences oligomer assembly and organization. Because cysteine oxidation can alter intermolecular interactions and promote disulfide bond formation, oxidation of Cys127 may shift the equilibrium among distinct oligomeric states, thereby affecting the ability of Hsp42 to sequester misfolded proteins and organize protein quality control compartments during oxidative stress. In this study, we identified a previously unreported Hsp42 trimer-like species that was detected using non-reducing SDS-PAGE from redox-challenged *trr1*Δ cells or WT cells treated with thiol-modifying agents (DVSF or H_2_O_2_), or recombinant Hsp42 purified from *E. coli*. Because Hsp42 contains only a single cysteine residue, a disulfide-linked dimer would be expected to form upon oxidation. Therefore, one possible explanation for the apparent trimer-like band was the presence of an additional interacting protein. However, IP-MS analysis showed that approximately 96% of the protein content within this band corresponded to Hsp42, strongly suggesting that it represents an Hsp42 homo-oligomer. Formation of this trimer-like species required both Cys127 oxidation and the N-terminal prion-like domain (PrLD). Interestingly, deletion of the PrLD converted the trimer-like species into an apparent dimer on non-reducing SDS-PAGE following Cys127 oxidation. These findings raise two possible scenarios. First, the trimer-like band may actually represent a disulfide-linked dimer whose electrophoretic mobility is altered by the presence of the PrLD. Alternatively, Cys127 oxidation and the PrLD may cooperate to stabilize a *bona fide* trimeric assembly that remains intact under SDS-PAGE conditions. To further characterize this species, we examined Hsp42 oligomers by BN-PAGE. However, no discrete trimeric Hsp42 complex was detected, consistent with previous reports showing only large Hsp42 oligomers under native conditions (39, 40). This observation suggests that the trimer-like species detected by SDS-PAGE is likely incorporated into larger oligomeric assemblies and becomes resolved only under SDS-denaturing conditions. Consistent with this model, both mutation of Cys127 to serine and treatment with reducing agents, increased the abundance of smaller Hsp42 oligomeric complexes. Together, these results indicate that disulfide bond formation at Cys127 promotes the assembly or stabilization of higher-molecular-weight Hsp42 oligomers during oxidative stress.

The PrLD is essential for Hsp42 foci formation during heat shock because it constitutes the primary substrate-binding region (38). Consistent with this role, we found that the PrLD is also required for Hsp42 foci formation in redox challenged *trr1*Δ cells, whereas deletion of the CTD had little effect and deletion of the ACD destabilized the protein. In contrast, deletion of the intrinsically disordered domain, which contains Cys127, or substitution of Cys127 with serine, only partially decreased foci formation in *trr1*Δ cells. These findings suggest that Cys127 is not a primary determinant of Hsp42 foci assembly under oxidative stress but instead serves a regulatory role. Notably, the IDD has previously been shown to modulate the dynamics and number of misfolded protein foci formed within a cell during heat shock and contains a regulatory phosphorylation site at Ser223. Phosphorylation of this residue disrupts Hsp42 co-localization with the disaggregase Hsp104 and impairs aggregate clearance and chaperone activity (46). Similarly, oxidation of Cys127 may represent an additional regulatory mechanism that modulates Hsp42 function during oxidative stress. Our data suggest that disulfide bond formation at Cys127 reinforces the stability of large Hsp42-containing assemblies that sequester unfolded/misfolded proteins under oxidative stress.

This conclusion is supported by the observation that C127 oxidation promotes the formation of higher-molecular-weight oligomeric species, whereas the C127S mutation reduces their stability and increases the proportion of Hsp42 present in the supernatant fraction of *trr1*Δ cells. Consistent with this interpretation, recombinant Hsp42 expressed in *E. coli* requires reducing agents to improve protein solubility and refolding efficiency (47). Moreover, we consistently obtained substantially higher yields of soluble Hsp42-C127S-6xHis than Hsp42-6xHis during purification from *E. coli* (Supplementary Figure S1).

To date, no study has comprehensively identified Hsp42 client proteins *in vivo* using Hsp42 as the affinity bait. In this work, we performed GFP-Trap pull-downs of Hsp42-GFP followed by mass spectrometry and identified 87 Hsp42-associated proteins in WT cells and 173 Hsp42-associated proteins in *trr1*Δ cells, with only eight proteins shared between the two datasets. The substantial increase in Hsp42-associated proteins in *trr1*Δ cells suggests that loss of *TRR1* imposes proteotoxic stress, leading to the accumulation of unfolded or misfolded proteins that are subsequently sequestered by Hsp42. In addition, disruption of cellular redox homeostasis in *trr1*Δ cells has been shown to induce ER stress and activate the unfolded protein response (UPR)(28, 30), which may further contribute to the increased burden of misfolded proteins and expansion of the Hsp42 interactome. Among the proteins associated with Hsp42 in WT cells were several proteasome subunits and the Hsp70 co-chaperone Sis1 (Supplementary Table S3), consistent with the established role of Hsp42 in directing protein quality-control substrates toward either refolding or degradation pathways. Remarkably, the Hsp42 interactome was almost completely remodeled in *trr1*Δ cells. Gene ontology analysis revealed a significant enrichment of mitochondrial proteins (59/173) and oxidoreductases (22/173) among Hsp42-associated proteins in *trr1*Δ cells. These findings suggest that mitochondrial and redox-related proteins are particularly susceptible to proteotoxic damage under oxidative stress and become preferentially sequestered by Hsp42. One possible explanation is that these proteins contain functionally important cysteine and methionine residues that act as redox-sensitive switches. Oxidation of these residues may alter protein conformation and stability, leading to protein unfolding or misfolding and subsequent sequestration by Hsp42. This effect may be especially pronounced in redox-related proteins, which frequently rely on such residues for their activity and regulation.

Interestingly, approximately 87 proteins identified in the Hsp42-C127S-GFP interactome overlapped with those found in Hsp42-GFP in *trr1*Δ cells, whereas an additional 140 proteins were uniquely associated with Hsp42-C127S-GFP. As a result, the enrichment of mitochondrial proteins and oxidoreductases observed in *trr1*Δ cells was no longer apparent in gene ontology analysis, despite only modest changes in the absolute numbers of proteins belonging to these categories. One possible explanation is that the C127S mutation promotes the formation of smaller Hsp42 oligomers, thereby increasing substrate accessibility and broadening the range of proteins that can associate with Hsp42. Consequently, the interactome becomes more diverse and less selectively enriched for mitochondrial and redox-related proteins.

Notably, most mitochondrial proteins identified in the Hsp42-GFP interactome of *trr1*Δ cells corresponded to mitochondrial precursor forms. Among these, the mitochondrial superoxide dismutase Sod2 was one of the most highly enriched proteins in both Hsp42-GFP and Hsp42-C127S-GFP pull-downs. Because Hsp42 is predominantly localized in the cytosol, the association of mitochondrial precursor proteins with Hsp42 suggests that mitochondrial protein import may be compromised under oxidative stress, resulting in the accumulation of import-defective precursor proteins in the cytosol. This interpretation is supported by a recent study showing that inhibition of mitochondrial protein import, either through clogging of import channels or during growth on non-fermentable carbon sources, promotes the formation of MitoStores, specialized cytosolic compartments containing Hsp42, Hsp104, and misfolded mitochondrial precursor proteins (48). It is therefore possible that disruption of cellular redox homeostasis in *trr1*Δ cells impairs mitochondrial protein import, leading to the accumulation of mitochondrial precursor proteins that are subsequently sequestered by Hsp42. Although we did not directly examine mitochondrial import efficiency or activation of the mitochondrial unfolded protein response (UPRmt), our findings support a model wherein oxidative stress caused by *TRR1* deletion affects both ER and mitochondrial proteostasis pathways, necessitating enhanced sequestration by Hsp42. This concept is further supported by the observation that *trr1Δ hsp42Δ* cells exhibit pronounced growth retardation compared to *trr1Δ* alone (19).

In summary, our data demonstrate that Cys127 of Hsp42 functions as a redox-sensitive residue that undergoes intermolecular disulfide bond formation under oxidative stress. Oxidation of Cys127 promotes the assembly of higher-order Hsp42 oligomers and contributes to the formation and stability of Hsp42-containing protein foci. These changes are accompanied by a marked remodeling of the Hsp42 interactome in *trr1*Δ cells, with mitochondrial proteins and oxidoreductase enzymes emerging as prominent classes of Hsp42-associated proteins. Together, our findings reveal a previously unrecognized mechanism by which redox regulation modulates Hsp42 function and proteostasis during oxidative stress. Future work will be needed to define the structural basis of Cys127-mediated oligomer remodeling and to determine how impaired thioredoxin-dependent redox control influences mitochondrial protein import and cytosolic quality-control pathways.

## Materials and Methods

### Strains and Plasmids

The yeast strains used in this study were derived from the BY4741 parental strain and are listed in Supplementary Table S1. The Hsp42-C127S mutant was generated using the pop-in/pop-out two-step allele replacement method with *Msc*I-digested pRS306-HSP42-C127S plasmid and selection on 5-FOA-containing medium. *TRR1* knockout strains were constructed by transforming PCR amplicons containing either the *URA3* or *LEU2* marker flanked by 50 bp homologous to the upstream and downstream regions of the *TRR1* coding sequence. Hsp42-GFP domain-deletion strains were generated by PCR amplification of the corresponding constructs containing the *HIS3MX6* cassette from the plasmids listed in Supplementary Table S1. The amplicons were flanked by 50 bp homologous to the regions upstream of the *HSP42* start codon and downstream of the *HSP42* stop codon, allowing replacement of the endogenous *HSP42* coding sequence. Correct integration and the resulting endogenous *HSP42-GFP* domain-deletion sequences were verified by PCR and DNA sequencing. The Hsp42-FLAG strain was generated using an extended forward primer containing the FLAG-tag coding sequence to amplify the *HIS3MX6* cassette, which was targeted to the 3′ end of the *HSP42* coding sequence. The strain expressing GFP under the control of the *HSP42* promoter was constructed by integrating *Van91*I-digested plasmid pIAU-HSP42prom-GFP into the *ADE2* locus. Yeast transformations were performed using the rapid yeast transformation protocol (49).

The plasmid pRS306-HSP42-C127S was constructed by PCR-based site-directed mutagenesis to generate DNA fragments containing the C127S mutation, which were subsequently assembled into *Sac*I- and *Kpn*I-digested pRS306 using NEBuilder® HiFi DNA Assembly Master Mix (NEB, Cat# E2621). The plasmid pRS426-BTN2-GFP was constructed by amplifying the BTN2-GFP sequence from genomic DNA of the yeast BTN2-GFP strain and assembling it into *Sac*I- and *Sal*I-digested pRS426 using NEBuilder® HiFi DNA Assembly Master Mix (NEB, Cat# E2621). Hsp42-GFP domain-deletion plasmids were generated using DNA fragments amplified from *Hsp42-GFP-HIS3MX6* or *Hsp42-C127S-GFP-HIS3MX6* genomic DNA templates. DNA fragments corresponding to the regions upstream and downstream of the deleted domain were assembled into *Sac*I- and *Sal*I-digested pRS416 using NEBuilder® HiFi DNA Assembly Master Mix (NEB, Cat# E2621). The C48S and C70S mutations were subsequently introduced into the *GFP* sequence of these domain-deletion plasmids by PCR-based mutagenesis and DNA assembly using NEBuilder® HiFi DNA Assembly Master Mix (NEB, Cat# E2621), generating a new set of Hsp42-GFP(C48S,C70S) domain-deletion plasmids. The integrating plasmid pIAU-HSP42prom-GFP was generated using *Sac*I-and *Spe*I-digested pRH2081 (50) as the backbone and assembled with the *HSP42* promoter fragment (500 bp upstream of the ATG start codon) and the *GFP-ADH1* terminator fragment using NEBuilder® HiFi DNA Assembly Master Mix (NEB, Cat# E2621). Plasmids pET28a-HSP42-6xHis, pET28a-HSP42-C127S-6xHis, and pET28a-HSP42-ΔPrLD-6xHis were constructed by assembling the *HSP42*, *HSP42-C127S*, or *HSP42-ΔPrLD* PCR amplicons, respectively, into *Nco*I- and *Not*I-digested pET28a using NEBuilder® HiFi DNA Assembly Master Mix (NEB, Cat# E2621). All plasmid constructs were verified by DNA sequencing and are listed in Supplementary Table S1.

### Yeast cultures

Yeast cells were cultured at 30°C with shaking to mid-logarithmic phase (OD_600_ ≈ 0.5–0.7) in standard YPD medium (1% yeast extract, 2% peptone, and 2% glucose) or in selective synthetic complete medium (SC; 2% glucose) lacking the appropriate amino acids for marker selection (Sunrise Science). For DVSF treatment, DVSF (≥96%, Sigma-Aldrich, Cat# V3700) was diluted in DMSO and added to mid-log phase cells at the indicated concentrations for 1 hour prior to harvesting. For H_2_O_2_ treatment, H_2_O_2_ (30%, Sigma-Aldrich, Cat# 216763) was diluted in water and added to mid-log phase cells at the indicated concentrations for the indicated times prior to harvesting.

### Immunoblotting

Proteins were isolated using glass bead lysis as previously described in (51). Whole-cell lysates were resuspended in 2x SDS sample buffer with or without reducing agent (5% β-mercaptoethanol) and boiled at 95°C for 5 min prior to separation on 4–12% SDS-PAGE gels (Thermo Fisher Scientific, Cat# XP04120BOX). Proteins were then transferred onto PVDF membranes (MilliporeSigma, Cat# IPVH00010) using a Mini Trans-Blot® Cell (Bio-Rad, Cat# 1703930). Membranes were blocked in 5% nonfat dry milk and incubated with primary antibodies followed by HRP-conjugated anti-mouse secondary antibody (R&D Systems, Cat# HAF007). Signals were detected using WesternBright ECL Spray (Advansta Inc., Cat# K-12049-D50) and visualized with a Bio-Rad ChemiDoc™ MP Imaging System. Primary antibodies used in this study included monoclonal anti-GFP (Roche, Cat# 11814460001), monoclonal anti-FLAG M2 (Sigma-Aldrich, Cat# F3165) and monoclonal anti-PGK1 (Invitrogen, Cat# 459250). Pgk1 was used as a loading control.

### Fluorescence microscopy

Yeast cells were wet-mounted on microscope slides and imaged immediately using an Olympus IX81-ZDC inverted fluorescence microscope equipped with a 100x oil-immersion objective lens and appropriate standard filter sets. Images were acquired using a Hamamatsu ORCA camera. For each biological replicate, foci quantification was performed by counting at least 100 cells across a minimum of three fields of view. The percentage of cells containing protein foci was calculated by dividing the number of cells with visible foci by the total number of cells counted. Statistical significance was assessed using Student’s t-test, with *p*-value < 0.05 considered significant.

### Protein purification and blue native PAGE

The recombinant proteins Hsp42-6xHis, Hsp42-C127S-6xHis, and Hsp42-ΔPrLD-6xHis were expressed in *E. coli* BL21(DE3) cells harboring the plasmids pET28a-HSP42-6xHis, pET28a-HSP42-C127S-6xHis, and pET28a-HSP42-ΔPrLD-6xHis, respectively. Bacterial cultures were grown in 300 mL LB medium supplemented with 50 μg/mL kanamycin and incubated at 37°C with shaking at 200 rpm. Protein expression was induced with 0.5 mM IPTG (Thermo Fisher Scientific, Cat# 15529019) when the cultures reached an OD_600_ of ∼0.5, followed by incubation at 30°C with shaking at 200 rpm for an additional 3 h before harvesting by centrifugation (7,000 rpm, RT, 10 min). Cell pellets were washed with cold 1x PBS and stored at −80°C until further use. Frozen cell pellets were resuspended in lysis buffer (50 mM Tris–HCl pH 7.5, 150 mM NaCl, 5% glycerol, 2% Triton X-100, 1 mM PMSF, 1x Halt EDTA-free protease inhibitor cocktail [Sigma, Cat#78425], and 20 mM imidazole) at a ratio of 150 OD units of cells per 15 mL lysis buffer, and disrupted by sonication using a QSONICA sonicator. The crude extracts (15 mL) were clarified by centrifugation (13,000 rpm, 4°C, 10 min) prior to incubation with 2 mL HisPur™ Cobalt Resin (1 mL gel bed volume; Thermo Fisher Scientific, Cat# 89965) at 4°C for 30 min. Unbound proteins were removed by washing with lysis buffer containing 50 mM imidazole, and the target proteins were eluted using elution buffer (50 mM Tris–HCl pH 7.5, 150 mM NaCl, 5% glycerol, and 500 mM imidazole). Protein purity was assessed by SDS-PAGE followed by Coomassie staining, and protein concentrations were determined using the Bradford protein assay (Thermo Fisher Scientific, Cat# 1856209).

Purified Hsp42-6xHis and Hsp42-C127S-6xHis proteins (3 µg each) were mixed with NativePAGE™ Sample Buffer (Thermo Fisher Scientific, Cat# BN2003) in the presence or absence of 10 mM DTT and loaded onto NativePAGE™ 4–16% Bis-Tris Gels (Thermo Fisher Scientific, Cat# BN1002BOX) together with NativeMark™ Unstained Protein Standard (Thermo Fisher Scientific, Cat# LC0725) as native protein markers. The anode buffer consisted of NativePAGE™ Running Buffer (Thermo Fisher Scientific, Cat# BN2001), while the cathode buffer was prepared by mixing NativePAGE™ Running Buffer with NativePAGE™ Cathode Additive (Thermo Fisher Scientific, Cat# BN2002) according to the manufacturer’s instructions. Samples were electrophoresed at 4°C at a constant voltage of 150 V for 60 minutes, followed by 250 V for the remainder of the run. Proteins were visualized using the Pierce™ Silver Stain Kit (Thermo Fisher Scientific, Cat# 24612).

### Cell fractionation

Cell lysates were prepared from mid-log phase cells using glass bead and lysis buffer containing 50 mM Tris-HCl (pH 7.5), 150 mM NaCl, 0.5 mM EDTA, 1 mM PMSF, and 1x Protease Inhibitor Cocktail (Sigma-Aldrich, Cat# P8215). Lysates were centrifuged at 3,500 rpm for 5 min at 4°C to remove cell debris. The protein concentration of the resulting supernatants was determined using the Bradford protein assay (Thermo Fisher Scientific, Cat# 1856209), and samples were adjusted to a final volume of 400 µl at a final protein concentration of 2 mg/ml using lysis buffer. Protein solutions were subsequently centrifuged at 15,000 rpm for 15 min at 4°C. The resulting supernatants were collected, while the pellets were resuspended in an equal volume of lysis buffer. Total, supernatant, and pellet fractions were analyzed by reducing SDS-PAGE followed by western blotting.

### Immunoprecipitation, mPEG-maleimide conjugation, and mass spectrometry

Mid-log phase cells were collected and resuspended in lysis buffer containing 50 mM Tris-HCl (pH 7.5), 0.5 mM EDTA 150 mM NaCl, 0.1% NP-40, 1 mM PMSF, and 1x Protease Inhibitor Cocktail (Sigma-Aldrich, Cat# P8215), followed by cell lysis using glass bead beating. Cell extracts were centrifuged at 3,500 rpm for 5 min at 4°C to remove cell debris, and the supernatants were collected. Protein concentrations were determined using the Bradford protein assay (Thermo Fisher Scientific, Cat# 1856209). A total of 1 mg of protein from each sample was used for immunoprecipitation. For Hsp42-FLAG or Hsp42-C127S-FLAG-containing cell lysates, 700 µL of lysate (1 mg total protein) was incubated with 40 µL anti-FLAG® M2 Magnetic Beads (20 µL bead bed volume; MilliporeSigma, Cat# M8823) with end-over-end rotation at room temperature for 1 h. The beads were washed three times with 700 µL lysis buffer before proteins were eluted with 100 µL 3xFLAG peptide (200 ng/µL). Eluted samples were subjected to non-reducing SDS-PAGE followed by western blotting to confirm the presence of target proteins, including the Hsp42 trimer-like form. SDS-PAGE gels were subsequently stained with Coomassie Blue, and the Hsp42 trimer-like band was excised, subjected to in-gel tryptic digestion, and analyzed by liquid chromatography–tandem mass spectrometry (LC-MS/MS) for protein identification.

To detect redox-active cysteine, eluted proteins were incubated with 10 mM TCEP to reduce cysteine thiols. TCEP was removed by buffer exchange into 1x PBS using Zeba™ Spin Desalting Columns, 7K MWCO (Thermo Fisher Scientific, Cat# 89882). Proteins were then incubated with 1 mM 10 kDa mPEG-maleimide (MilliporeSigma, Cat# JKA3119) at room temperature for 30 min before being subjected to non-reducing SDS-PAGE followed by western blotting.

For GFP, Hsp42-GFP, or Hsp42-C127S-GFP containing cell lysates, 500 µL of lysate (1 mg total protein) was incubated with 40 µL ChromoTek GFP-Trap® Magnetic Particles M-270 (20 µL bead bed volume; Proteintech, Cat# gtd) at 4°C for 1 h with end-over-end rotation. The beads were washed four times with 500 µL lysis buffer and proteins were eluted with 100 µL 1x sample buffer at 70°C for 15 min. Beads were subsequently removed, and the eluted proteins were subjected to in-gel tryptic digestion followed by liquid chromatography–tandem mass spectrometry (LC-MS/MS) analysis for interactome identification. Protein abundances detected in immunoprecipitates were compared directly across samples, with two biological replicates for each condition, to identify enriched interacting proteins. Differential analysis was performed using the moderated t-test implemented in the limma R package (52). Missing values were assumed to be missing not at random (MNAR) and were imputed by sampling from a downshifted normal distribution defined as N(μ − 1.8σ, 0.35σ), where μ and σ represent the mean and standard deviation of all quantified values, respectively. Multiple hypothesis testing correction was performed using the Benjamini–Hochberg procedure. Proteins exhibiting at least eightfold enrichment with a *p*-value < 0.05 were considered statistically significant. Gene ontology (GO) enrichment analyses of client proteins were performed using g:Profiler (53) to identify significantly enriched biological processes, molecular functions, and cellular components. Enrichment analysis was conducted against the *Saccharomyces cerevisiae* reference genome using the g:SCS algorithm with multiple testing correction. GO terms with an adjusted *p*-value < 0.05 were considered significantly enriched.

## Acknowledgements

This work was supported by grant R35GM149196 from the National Institutes of Health to K.A.M. We thank Dr. Anna Malovannaya, Antrix Jain, and Shirley Wang of the Mass Spectrometry Proteomics Core at Baylor College of Medicine for their excellent technical support. The BCM Mass Spectrometry Proteomics Core (RRID:SCR_027015) is supported by the Dan L. Duncan Comprehensive Cancer Center NIH award (P30 CA125123), CPRIT Core Facility Award (RP210227). We also thank Dr. Sheng Pan, Dr. Lakmini Senavirathna, and Li Li of the Clinical and Translational Proteomics Service Center at UTHealth Houston for their excellent technical support. We thank Dr. James West (College of Wooster) for helpful discussions.

## Data Availability

The mass spectrometry proteomics data have been deposited to the ProteomeXchange Consortium via the PRIDE partner repository with the dataset identifier PXD080899.

## Abbreviations

sHSP: small heat shock protein
DVSF: divinyl sulfone
ACD: α-crystallin domain
NTD: N-terminal domain
CTD: C-terminal domain
IDD: intrinsic disorder domain
PrLD: prion-like domain
mPEG-MAL: methoxy polyethylene glycol maleimide
BN-PAGE: blue native polyacrylamide gel electrophoresis
ROS: reactive oxygen species.

## Supplementary Figure Legends

**Figure S1.**
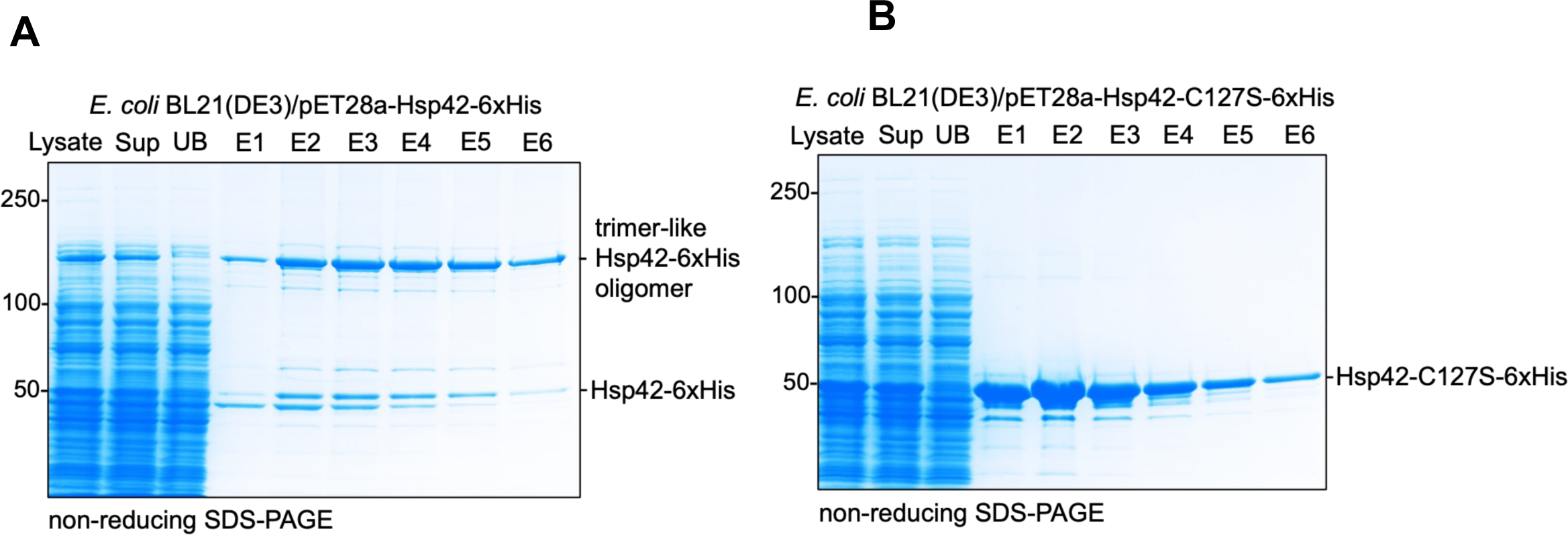
Hsp42-6xHis and Hsp42-C127S-6xHis purifications from *E. coli* cells. Protein purification was performed as described in the Materials and Methods section. Sup, cell lysate supernatant; UB, unbound material; E1–E6, elution fractions.

**Figure S2.**
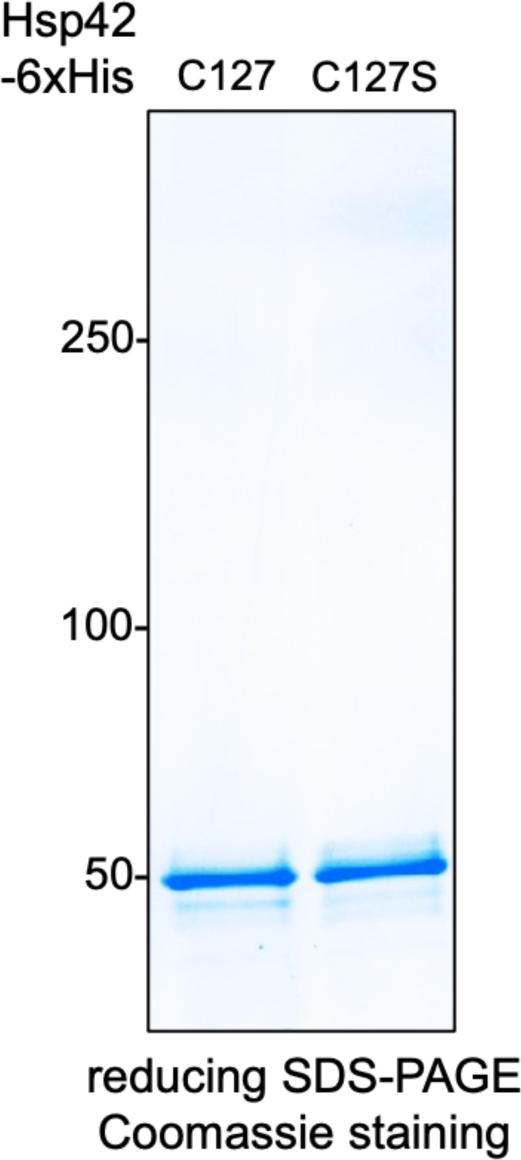
**The comparable amounts of purified Hsp42-6xHis and Hsp42-C127S-6xHis used for BN-PAGE analysis were verified by reducing SDS-PAGE followed by Coomassie staining.**

**Figure S3.**
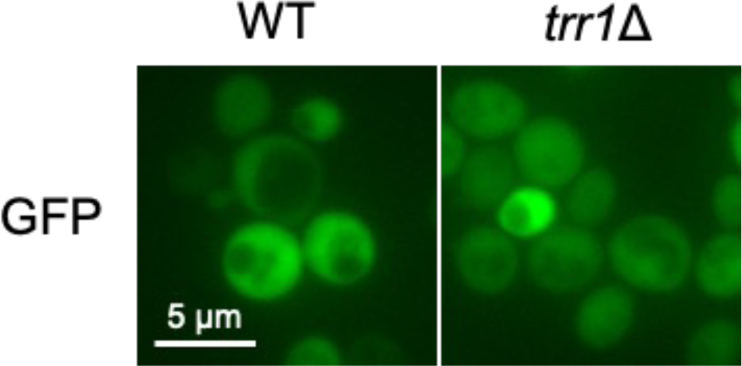
**Endogenously expressed GFP under the *HSP42* promoter exhibits diffuse localization in WT and *trr1*Δ mid-log phase cells.**

