## Supplementary Table and Data for "Intermolecular disulfide bond formation promotes Hsp42 higher-order assembly and shapes client selection in yeast"

**Precursor Mitochondrial Proteins Identified by IP–MS with MTS-Containing Peptides**

MTSs are highlighted in red.

**1. Aim17**

**MLRSNLCRGSRILARLTTTPRT**YTSAATAAAANRGHIIKTYFNRDSTTITFSMEESSKPVSVCFNNVFLRDASHSAKLVTTGE
LYHNEKLTAPQDIQISEDGKSLVVKWKDGGHHQFPLQFFIDYKGSSFVSPATRKQESRYRPQLWNKRILKDNVKDLLSVSYN
EFIDPKDDSKLFQTLVNLQKFGIAFISGTPSSSSEGLTIQKICERIGPIRSTVHGEGTFDVNASQATSVNAHYANKDLPLHTDL
PFLENVPGFQILQSLPATEGEDPNTRPMNYFVDAFYATRNVRESDFEAYEALQIVPVNYIYENGDKRYYQSKPLIEHHDINE
DNTLLGNYEALIKCINYSPPYQAPFTFGIYDKPSDLNNNLDLNLITTPAKLTERFLFKSFIRGLNLFESHINDFNNQFRLQLPEN
CCVIFNNRRILHANSLTSSNQQWLKGCYFDSDTFKSKLKFLEEKFPHDK*

Peptides detected in IP-MS

| WT Hsp42-GFP | *trr1*Δ Hsp42-GFP | *trr1*Δ Hsp42-C127S-GFP |
|---|---|---|
| LTAPQDIQISEDGK | LTAPQDIQISEDGK<br>LFQTLVNLQK<br>LTAPQDIQISEDGK<br>**T**YTSAATAAAANR | LTAPQDIQISEDGK |

### 2. Ald4

**MFSRSTLCLKTSASSIGRLQLRY**FSHLPMTVPIKLPNGLEYEQPTGLFINNKFVPSKQNKTFEVINPSTEEEICHIYEGREDD
VEEAVQAADRAFSNGSWNGIDPIDRGKALYRLAELIEQDKDVIASIETLDNGKAISSSRGDVDLVINYLKSSAGFADKIDGRMI
DTGRTHFSYTKRQPLGVCGQIIPWNFPLLMWAWKIAPALVTGNTVVLKTAESTPLSALYVSKYIPQAGIPPGVINIVSGFGKIV
GEAITNHPKIKKVAFTGSTATGRHIYQSAAAGLKKVTLELGGKSPNIVFADAELKKAVQNIILGIYYNSGEVCCAGSRVYVEES
IYDKFIEEFKAASESIKVGDPFDESTFQGAQTSQMQLNKILKYVDIGKNEGATLITGGERLGSKGYFIKPTVFGDVKEDMRIV
KEEIFGPVVTVTKFKSADEVINMANDSEYGLAAGIHTSNINTALKVADRVNAGTVWINTYNDFHHAVPFGGFNASGLGREMS
VDALQNYLQVKAVRAKLDE*

Peptides detected in IP-MS

| WT Hsp42-GFP | *trr1*Δ Hsp42-GFP | *trr1*Δ Hsp42-C127S-GFP |
| --- | --- | --- |
| EMSVDALQNYLQVK<br>IAPALVTGNTVVLK<br>KVTLELGGK<br>LPNGLEYEQPTGLFINN<br>K<br>NEGATLITGGER<br>TAESTPLSALYVSK<br>VGDPFDESTFQGAQTS<br>QMQLNK<br>VTLELGGK | AFSNGSWNGIDPIDR<br>DVIASIETLDNGK<br>EEIFGPVVTVTK<br>EMSVDALQNYLQVK<br>GDVDLVINYLK<br>GYFIKPTVFGDVK<br>HIYQSAAAGLK<br>IAPALVTGNTVVLK<br>IVGEAITNHPK<br>KVAFTGSTATGR<br>KVTLELGGK<br>LAELIEQDK<br>LPNGLEYEQPTGLFINNK<br>NEGATLITGGER<br>PTVFGDVK<br>SADEVINMANDSEYGLAAGIHT<br>SNINTALK<br>SPNIVFADAELK<br>SPNIVFADAELKK<br>SSAGFADK | EMSVDALQNYLQVK<br>HIYQSAAAGLK<br>IAPALVTGNTVVLK<br>LAELIEQDK<br>LPNGLEYEQPTGLFINNK<br>NEGATLITGGER<br>SADEVINMANDSEYGLAAGIHT<br>SNINTALK<br>SPNIVFADAELK<br>SPNIVFADAELKK<br>TAESTPLSALYVSK<br>VAFTGSTATGR<br>VGDPFDESTFQGAQTSQMQL<br>NK<br>VTLELGGK |

| | |
|---|---|
| | TAESTPLSALYVSK<br>VAFTGSTATGR<br>VGDPFDESTFQGAQTSQMQL<br>NK<br>VTLELGGK<br>VYVEESIYDK<br>**Y**FSHLPMTVPIK |

**3. Bat1**

MLQRHSLKLGKFSIRTLATGAPLDASKLKITRNPNPSKPRPNEELVFGQTFTDHMLTIPWSAKEGWGTPHIKPYGNLSLDPS
ACVFHYAFELFEGLKAYRTPQNTITMFRPDKNMARMNKSAARICLPTFESEELIKLTGKLIEQDKHLVPQGNGYSLYIRPTMI
GTSKGLGVGTPSEALLYVITSPVGPYYKTGFKAVRLEATDYATRAWPGGVGDKKLGANYAPCILPQLQAAKRGYQQNLWLF
GPEKNITEVGTMNVFFVFLNKVTGKKELVTAPLDGTILEGVTRDSVLTLARDKLDPQEWDINERYYTITEVATRAKQGELLEA
FGSGTAAVVSPIKEIGWNNEDIHVPLLPGEQCGALTKQVAQWIADIQYGRVNYGNWSKTVADLN*

Peptides detected in IP-MS

| WT Hsp42-GFP | *trr1*Δ Hsp42-GFP | *trr1*Δ Hsp42-C127S-GFP |
| --- | --- | --- |
| DSVLTLAR<br>ELVTAPLDGTILEGVTR<br>**T**LATGAPLDASK<br>YYTITEVATR | DSVLTLAR<br>ELVTAPLDGTILEGVTR<br>LEATDYATR<br>QGELLEAFGSGTAAVVSPIK<br>**T**LATGAPLDASK<br>YYTITEVATR | ELVTAPLDGTILEGVTR<br>LEATDYATR |

**4. Cbp6**

MSSSQVVRDSAKKLVNLLEKYPKDRIHHLVSFRDVQIARFRRVAGLPNVDDKGKSIKEKKPSLDEIKSIINRTSGPLGLNKEM
LTKIQNKMVDEKFTEESINEQIRALSTIMNNKFRNYYDIGDKLYKPAGNPQYYQRLINAVDGKKKESLFTAMRTVLFGK*

Peptides detected in IP-MS

| WT Hsp42-GFP | *trr1*Δ Hsp42-GFP | *trr1*Δ Hsp42-C127S-GFP |
|---|---|---|
| FTEESINEQIR<br>TSGPLGLNK | ALSTIMNNK<br>EMLTKIQNK<br>ESLFTAMR<br>FTEESINEQIR<br>LVNLLEK<br>NYYDIGDK<br>PAGNPQYYQR<br>TSGPLGLNK | ALSTIMNNK<br>FTEESINEQIR<br>LVNLLEK<br>PAGNPQYYQR<br>TSGPLGLNK<br>VAGLPNVDDK |

**5. Gcv2**

<u>MLRTRVTALLCRATVRSSTNYVSLARTRS</u>FHSQSILLKTAATDITSTQYSRIFNPDLKNIDRPLDTFARRHLGPSPSDVKKML
KTMGYSDLNAFIEELVPPNILKRRPLKLEAPSKGFCEQEMLQHLEKIANKNHYKVKNFIGKGYYGTILPPVIQRNLLESPEWY
TSYTPYQPEISQGRLEALLNFQTVVSDLTGLPVANASLLDEGTAAGEAMLLSFNISRKKKLKYVIDKKLHQQTKSVLHTRAKP
FNIEIIEVDCSDIKKAVDVLKNPDVSGCLVQYPATDGSILPPDSMKQLSDALHSHKSLLSVASDLMALTLLKPPAHYGADIVLG
SSQRFGVPMGYGGPHAAFFAVIDKLNRKIPGRIVGISKDRLGKTALRLALQTREQHIKRDKATSNICTAQALLANVASSYCVY
HGPKGLQNISRRIFSLTSILANAIENDSCPHELINKTWFDTLTIKLGNGISSEQLLDKALKEFNINLFAVDTTTISLALDETTTKA
DVENLLKVFDIENSSQFLSEDYSNSFPREFQRTDEILRNEVFHMHHSETAMLRYLHRLQSRDLSLANSMIPLGSCTMKLNS
TVEMMPITWPQFSNIHPFQPSNQVQGYKELITSLEKDLCSITGFDGISLQPNSGAQGEYTGLRVIRSYLESKGENHRNVCLI
PVSAHGTNPASAAMAGLKVVPVNCLQDGSLDLVDLKNKAEQHSKELAAVMITYPSTYGLFEPGIQHAIDIVHSFGGQVYLD
GANMNAQVGLTSPGDLGADVCHLNLHKTFSIPHGGGGPAGAPICVKSHLIPHLPKHDVVDMITGIGGSKSIDSVSSAPYGN
ALVLPISYAYIKMMGNEGLPFSSVIAMLNSNYMMTRLKDHYKILFVNEMSTLKHCAHEFIVDLREYKAKGVEAIDVAKRLQDY
GFHAPTLAFPVPGTLMIEPTESENLEELDRFCDAMISIKEEINALVAGQPKGQILKNAPHSLEDLITSSNWDTRGYTREEAAY
PLPFLRYNKFWPTVARLDDTYGDMNLICTCPSVEEIANETE*

Peptides detected in IP-MS

| WT Hsp42-GFP | *trr1*Δ Hsp42-GFP | *trr1*Δ Hsp42-C127S-GFP |
|---|---|---|
| TAATDITSTQYSR | GVEAIDVAK<br>GYYGTILPPVIQR<br>IFNPDLK<br>LGNGISSEQLLDK<br><u>SSTNYVSLAR</u><br>TAATDITSTQYSR | TAATDITSTQYSR |

**6. Mam33**

MFLRSVNRAVTRSILTTPKPAVVKSSWRVFTVANSKRCFTPAAIMRNQETQRVGDILQSELKIEKETLPESTSLDSFNDFLNK
YKFSLVETPGKNEAEIVRRTESGETVHVFFDVAQIANLPYNNAMDENTEQNEDGINEDDFDALSDNFANVNVVISKESASEP
AVSFELLMNLQEGSFYVDSATPYPSVDAALNQSAEAEITRELVYHGPPFSNLDEELQESLEAYLESRGVNEELASFISAYSE
FKENNEYISWLEKMKKFFH*

Peptides detected in IP-MS

| WT Hsp42-GFP | *trr1*$\Delta$ Hsp42-GFP | *trr1*$\Delta$ Hsp42-C127S-GFP |
|---|---|---|
| ETLPESTSLDSFNDFLNK<br>FSLVETPGKNEAEIVR<br>VGDILQSELK | ETLPESTSLDSFNDFLNK<br>FSLVETPGK<br>FSLVETPGKNEAEIVR<br>SILTTPK<br>VFTVANSK<br>VGDILQSELK | FSLVETPGK<br>FSLVETPGKNEAEIVR<br>SILTTPK |

**7. Mas1**

MFSRTASKFRNTRRLLSTISSQIPGTRTSKLPNGLTIATEYIPNTSSATVGIFVDAGSRAENVKNNGTAHFLEHLAFKGTQNR
SQQGIELEIENIGSHLNAYTSRENTVYYAKSLQEDIPKAVDILSDILTKSVLDNSAIERERDVIIRESEEVDKMYDEVVFDHLHEI
TYKDQPLGRTILGPIKNIKSITRTDLKDYITKNYKGDRMVLAGAGAVDHEKLVQYAQKYFGHVPKSESPVPLGSPRGPLPVF
CRGERFIKENTLPTTHIAIALEGVSWSAPDYFVALATQAIVGNWDRAIGTGTNSPSPLAVAASQNGSLANSYMSFSTSYADS
GLWGMYIVTDSNEHNVQLIVNEILKEWKRIKSGKISDAEVNRAKAQLKAALLLSLDGSTAIVEDIGRQVVTTGKRLSPEEVFE
QVDKITKDDIIMWANYRLQNKPVSMVALGNTSTVPNVSYIEEKLNQ*

Peptides detected in IP-MS

| WT Hsp42-GFP | *trr1*Δ Hsp42-GFP | *trr1*Δ Hsp42-C127S-GFP |
|---|---|---|
| LQNKPVSMVALGNTSTVPNVSYIEEK | AVDILSDILTK<br>LLSTISSQIPGTR<br>LQNKPVSMVALGNTSTVPNVSYIEEK<br>LSPEEVFEQVDK<br>MVLAGAGAVDHEK<br>SESPVPLGSPR<br>SLQEDIPK<br>SVLDNSAIER | AVDILSDILTK<br>LQNKPVSMVALGNTSTVPNVSYIEEK<br>MVLAGAGAVDHEK<br>SESPVPLGSPR<br>SVLDNSAIER |

**8. Mmf1**

MFLRNSVLRTAPVLRRGITTLTPVSTKLAPPAAASYSQAMKANNFVYVSGQIPYTPDNKPVQGSISEKAEQVFQNVKNILAE
SNSSLDNIVKVNVFLADMKNFAEFNSVYAKHFHTHKPARSCVGVASLPLNVDLEMEVIAVEKN*

Peptides detected in IP-MS

| WT Hsp42-GFP | *trr1*Δ Hsp42-GFP | *trr1*Δ Hsp42-C127S-GFP |
|---|---|---|
| AEQVFQNVK<br>**G**ITTLTPVSTK<br>NILAESNSSLDNIVK<br>VNVFLADMK | AEQVFQNVK<br>ANNFVYVSGQIPYTPDNKPVQGSISEK<br>**G**ITTLTPVSTK<br>LAPPAAASYSQAMK<br>NILAESNSSLDNIVK<br>VNVFLADMK | **G**ITTLTPVSTK<br>NILAESNSSLDNIVK<br>VNVFLADMK |

**9. Mrpl44**

MITKYFSKVIVRFNPFGKEAKVARLVLAAIPPTQRNMGTQIQSEIISDYNKVKPLVKVTYKDKKEMEVDPSNMNFQELANHFD RHSKQLDLKHMLEMH*

Peptides detected in IP-MS

| WT Hsp42-GFP | *trr1*Δ Hsp42-GFP | *trr1*Δ Hsp42-C127S-GFP |
|---|---|---|
| | EMEVDPSNMNFQELANHFDR<br>LVLAAIPPTQR<br>**N**MGTQIQSEIISDYNK | **N**MGTQIQSEIISDYNK |

**10. Mtg1**

MHINVRGTRKIISNVSSFTPRYEFPKYSMPLTDFKGHQVKALKTFEKLLPQMNMIIELRDIRAPLSTRNVVFDRIARKEHDVM
KLVVYTRKDLMPGNKPYIGKLKNWHEELGEKFILLDCRNKTDVRNLLKILEWQNYELETNGGYLPMGYRALITGMPNVGKS
TLINSLRTIFHNQVNMGRKFKKVAKTGAEAGVTRATSEVIRVTSRNTESRNEIYLIDTPGIGVPGRVSDHNRMLGLALCGSVK
NNLVDPIFQADYLLYLMNLQNLNDGRTELYPGSTNSPTNDIYDVLRRLQVNKSQNEKSTAIEWTNKWRLHGKGIIFDPEVLL
NNDEFSYKNYVNDQLEKLGDLSYEGLSNKLKGNPNQVF*

Peptides detected in IP-MS

| WT Hsp42-GFP | *trr1*Δ Hsp42-GFP | *trr1*Δ Hsp42-C127S-GFP |
|---|---|---|
| | ALITGMPNVGK<br>IISNVSSFTPR<br>NEIYLIDTPGIGVPGR | ALITGMPNVGK<br>IISNVSSFTPR<br>NEIYLIDTPGIGVPGR |

**11. Pda1**

MLAASFKRQPSQLVRGLGAVLRTPTRIGHVRTMATLKTTDKKAPEDIEGSDTVQIELPESSFESYMLEPPDLSYETSKATLL
QMYKDMVIIRRMEMACDALYKAKKIRGFCHLSVGQEAIAVGIENAITKLDSIITSYRCHGFTFMRGASVKAVLAELMGRRAG
VSYGKGGSMHLYAPGFYGGNGIVGAQVPLGAGLAFAHQYKNEDACSFTLYGDGASNQGQVFESFNMAKLWNLPVVFCC
ENNKYGMGTAASRSSAMTEYFKRGQYIPGLKVNGMDILAVYQASKFAKDWCLSGKGPLVLEYETYRYGGHSMSDPGTTY
RTRDEIQHMRSKNDPIAGLKMHLIDLGIATEAEVKAYDKSARKYVDEQVELADAAPPPEAKLSILFEDVYVKGTETPTLRGRI
PEDTWDFKKQGFASRD*

Peptides detected in IP-MS

| WT Hsp42-GFP | *trr1*Δ Hsp42-GFP | *trr1*Δ Hsp42-C127S-GFP |
|---|---|---|
| ATLLQMYK | ATLLQMYK | ATLLQMYK |
| GPLVLEYETYR | AVLAELMGR | AVLAELMGR |
| GQYIPGLK | GPLVLEYETYR | GPLVLEYETYR |
| GTETPTLR | GQYIPGLK | GQYIPGLK |
| KYVDEQVELADAAPPPEAK | GRIPEDTWDFKK | KYVDEQVELADAAPPPEAK |
| LDSIITSYR | GTETPTLR | LDSIITSYR |
| SKNDPIAGLK | IPEDTWDFKK | LSILFEDVYVK |
| VNGMDILAVYQASK | KYVDEQVELADAAPPPEAK | NDPIAGLK |
| YGMGTAASR | LDSIITSYR | RAGVSYGK |
| YVDEQVELADAAPPPEAK | LSILFEDVYVK | RGQYIPGLK |
| | QPSQLVR | SSAMTEYFK |
| | SSAMTEYFK | SSAMTEYFKR |
| | VNGMDILAVYQASK | VNGMDILAVYQASK |
| | YGMGTAASR | YGGHSMSDPGTTYR |
| | YVDEQVELADAAPPPEAK | YGMGTAASR |
| | | YVDEQVELADAAPPPEAK |

**12. Psd1**

MSIMPVKNALAQGRTLLMGRMPAVKFSTRMQLRNRTAVLWNRKFSTRLFVQQRRSSGEIVDRAKAAAANSGRKQVSMKW
VVLTSFTIVLGTILLVSRNDSTEEDATEGKKGRRTRKIKIFNNNWLFFCYSTLPLNAMSRLWGQVNSLTLPIWVRPWGYRLYS
FLFGVNLDEMEDPDLTHYANLSEFFYRNIKPGTRPVAQGEDVIASPSDGKILQVGIINSETGEIEQVKGMTYSIKEFLGTHSH
PLMSKSASSLDLTSDEEKHREFARVNRIQLAGSEDTEQPLLNFKNEGDQSVREFKPSVSKNIHLLSQLSLNYFSNGFSCSE
PHDTELFFAVIYLAPGDYHHFHSPVDWVCKVRRHFPGDLFSVAPYFQRNFPNLFVLNERVALLGSWKYGFFSMTPVGATN
VGSIKLNFDQEFVTNSKSDKHLEPHTCYQAVYENASKILGGMPLVKGEEMGGFELGSTVVLCFEAPTEFKFDVRVGDKVK
MGQKLGIIGKNDLK*

Peptides detected in IP-MS

| WT Hsp42-GFP | *trr1*Δ Hsp42-GFP | *trr1*Δ Hsp42-C127S-GFP |
|---|---|---|
| IQLAGSEDTEQPLLNFK | ILGGMPLVK | ILGGMPLVK<br>ILQVGIINSETGEIEQVK<br>IQLAGSEDTEQPLLNFK<br>MSIMPVK |

**13. Ptc7**

MFANVGFRTLRVSRGPLYGSCSQIISFSKRTFYSSAKSGYQSNNSHGDAYSSGSQSGPFTYKTAVAFQPKDRDDLIYQKLK
DSIRSPTGEDNYFVTSNNVHDIFAGVADGVGGWAEHGYDSSAISRELCKKMDEISTALAENSSKETLLTPKKIIGAAYAKIRD
EKVVKVGGTTAIVAHFPSNGKLEVANLGDSWCGVFRDSKLVFQTKFQTVGFNAPYQLSIIPEEMLKEAERRGSKYILNTPRD
ADEYSFQLKKKDIIILATDGVTDNIATDDIELFLKDNAARTNDELQLLSQKFVDNVVSLSKDPNYPSVFAQEISKLTGKNYSGG
KEDDITVVVVRVD*

Peptides detected in IP-MS

| WT Hsp42-GFP | *trr1*Δ Hsp42-GFP | *trr1*Δ Hsp42-C127S-GFP |
|---|---|---|
| DPNYPSVFAQEISK<br>TNDELQLLSQK | DADEYSFQLK<br>DPNYPSVFAQEISK<br>FVDNVVSLSK<br>IIGAAYAK<br>MFANVGFR<br>TAVAFQPK | FVDNVVSLSK<br>DPNYPSVFAQEISK |

**14. Sod2**

MFAKTAAANLTKKGGLSLLSTTARRTKVTLPDLKWDFGALEPYISGQINELHYTKHHQTYVNGFNTAVDQFQELSDLLAKEP
SPANARKMIAIQQNIKFHGGGFTNHCLFWENLAPESQGGGEPPTGALAKAIDEQFGSLDELIKLTNTKLAGVQGSGWAFIVK
NLSNGGKLDVVQTYNQDTVTGPLVPLVAIDAWEHAYYLQYQNKKADYFKAIWNVVNWKEASRRFDAGKI*

Peptides detected in IP-MS

| WT Hsp42-GFP | *trr1*Δ Hsp42-GFP | *trr1*Δ Hsp42-C127S-GFP |
|---|---|---|
| MIAIQQNIK | AIDEQFGSLDELIK<br>EPSPANAR<br>GGLSLLSTTAR<br>GGLSLLSTTARR<br>KGGLSLLSTTAR<br>MIAIQQNIK<br>VTLPDLK | AIDEQFGSLDELIK<br>EPSPANAR<br>GGLSLLSTTAR<br>KGGLSLLSTTAR<br>MIAIQQNIK<br>VTLPDLK |

**15. YJL133C-A/ Dpi8**

MIAQSTRLAAAVSSSAASAGVSRIAASAMASTIFKRSPGNSFNSFKEYRENAKTYGPLSASLATRRHLAHAPKL*

Peptides detected in IP-MS

| WT Hsp42-GFP | *trr1*Δ Hsp42-GFP | *trr1*Δ Hsp42-C127S-GFP |
| --- | --- | --- |
| **L**AAAVSSSAASAGVSR | **L**AAAVSSSAASAGVSR | **L**AAAVSSSAASAGVSR |

**16. Yme2**

MLLVRTTSLNVSRMPVPCLARGIGILKGKYRLANLMNAQPSVRHVSSEIQQKDQQAGESNTATDTGVIHKSDEETLIYFDNV
YARTTSVWNPTLWYNLLLRNQSRDAVREKIRNLASPPNNPIYGLELKSTIPVKRDGGVFATFVVPPKYTKAQVNSLIQQNTA
RESSKNLLSYFTRASAFPVKGSPWIEDLRRLPSTTIVIKFQGPALTEEEIYSLFRRYGTIIDIFPPTAANNNVAKVRYRSFRGAI
SAKNCVSGIEIHNTVLHIQYENIRRGHLVSNFFTNHTRIAIPVLFALLSIFAVLVFDPIREFSIEQKITHKYSLSWDNKFWKQLKT
LTSSTMTSIKYYWGGPDDNHQRKHLWEERIEKVNDLKMWLEENNNTFVVIRGPRGSGKHDLVMQHTLQNRANVLYLDCD
KLIKSRTDPMFLKNAASQLGYFPIFPWIDSVTGVLDLTVQGLTGQKTGLSETKESRFRNMLTTSLMSIRRIALKNYKAFVSTG
DGTVNVKEEDYLQQHPEAKPVIVIDRFEGKSEINGFVYKELSDWAAMLVQMNIAHVIFLTETVASNQRLSESLPNQVFKNLIL
SDASKENSRNYVLSQLEDYLYYNKKSKGENVKEPESEKETAENNDSDSEADTSVKKAEVILNEKELQEIDASLEPLGGRML
DLQAFVRRVKSGEEPSEAVDKMIEQASEQITQMFLSDKIDSNKSAQAWELIELLSANPVIPFHEIVNKPLFKAAPETGIMELE
NNGLITVSRDRGVLQEIRPAKPLYRAAFTYLINDPELAKVLKTRYLLKVVGFETGRIKKWEEELKPLGKVPDQKLFKTRLDYL
SGKINASNAVITKCEEEIKNLSK*

Peptides detected in IP-MS

| WT Hsp42-GFP | *trr1*Δ Hsp42-GFP | *trr1*Δ Hsp42-C127S-GFP |
|---|---|---|
| DQQAGESNTATDTGVIHK | AAPETGIMELENNGLITVSR | AAPETGIMELENNGLITVSR |
| ELQEIDASLEPLGGR | AFVSTGDGTVNVK | AFVSTGDGTVNVK |
| | AQVNSLIQQNTAR | AQVNSLIQQNTAR |
| | DGGVFATFVVPPK | DGGVFATFVVPPK |
| | DQQAGESNTATDTGVIHK | DQQAGESNTATDTGVIHK |
| | DRGVLQEIRPAK | ELQEIDASLEPLGGR |
| | ELQEIDASLEPLGGR | ETAENNDSDSEADTSVK |
| | ETAENNDSDSEADTSVK | INASNAVITK |
| | INASNAVITK | IRNLASPPNNPIYGLELK |
| | LANLMNAQPSVR | MLDLQAFVR |
| | LPSTTIVIK | NLASPPNNPIYGLELK |
| | MLDLQAFVR | NLILSDASK |
| | NLASPPNNPIYGLELK | SDEETLIYFDNVYAR |
| | NLILSDASK | TLTSSTMTSIK |
| | SGEEPSEAVDK | |

| | |
|---|---|
| | TLTSSTMTSIK<br>TRLDYLSGK<br>VVGFETGR |

Table S1. Yeast strains and plasmids

| Strains | Description | Reference |
|---|---|---|
| BY4741 | *MATα his3Δ1 leu2Δ0 met15Δ0 ura3Δ0* | Laboratory stock |
| Btn2-GFP | BY4741 *HSP26-GFP::HIS3* | Yeast GFP clone collection, Thermo Fisher Scientific |
| Hsp26-GFP | BY4741 *HSP26-GFP::HIS3* | Yeast GFP clone collection, Thermo Fisher Scientific |
| Hsp42-GFP | BY4741 *HSP42-GFP::HIS3* | Yeast GFP clone collection, Thermo Fisher Scientific |
| Hsp42 C127S-GFP | BY4741 *HSP42-C127S-GFP::HIS3* | This study |
| *trr1Δ* Hsp26-GFP | BY4741 *HSP26-GFP::HIS3 trr1Δ::URA3* | (1) |
| *trr1Δ* Hsp42-GFP | BY4741 *HSP42-GFP::HIS3 trr1Δ::URA3* | (1) |
| *trr1Δ* Hsp42-C127S-GFP | BY4741 *HSP42-C127S-GFP::HIS3 trr1Δ::URA3* | This study |
| Hsp42-FLAG | BY4741 *HSP42-FLAG::HIS3* | This study |
| *trr1Δ* Hsp42-FLAG | BY4741 *HSP42-FLAG::HIS3 trr1Δ::LEU2* | This study |
| Hsp42-C127S-FLAG | BY4741 *HSP42-C127S-FLAG::HIS3* | This study |
| *trr1Δ* Hsp42-ΔNTD-GFP | BY4741 *HSP42-Δ(1-726)-GFP::HIS3 trr1Δ::LEU2* | This study |
| *trr1Δ* Hsp42-ΔPrLD-GFP | BY4741 *HSP42-Δ(1-297)-GFP::HIS3 trr1Δ::LEU2* | This study |
| *trr1Δ* Hsp42-ΔIDD-GFP | BY4741 *HSP42-Δ(298-726)-GFP::HIS3 trr1Δ::LEU2* | This study |
| *trr1Δ* Hsp42-ΔACD-GFP | BY4741 *HSP42-Δ(727-1041)-GFP::HIS3 trr1Δ::LEU2* | This study |
| *trr1Δ* Hsp42-ΔCTD-GFP | BY4741 *HSP42-Δ(1042-1125)-GFP::HIS3 trr1Δ::LEU2* | This study |
| HSP42prom-GFP | BY4741 *ADE2::HSP42prom(500)-GFP-ADH1ter-URA3* | This study |

| | | |
|---|---|---|
| *trr1*Δ HSP42prom-GFP | BY4741 *ADE2::HSP42prom(500)-GFP-ADH1ter-URA3 trr1Δ::LEU2* | This study |
| **Plasmids** | | |
| pRS306-HSP42-C127S | *HSP42(379–381:TGT → TCG)*; YIp, *URA3*, *AmpR* | This study |
| pRS426-BTN2-GFP | *BTN2-GFP*, 2μ, *URA3*, *AmpR* | This study |
| pRS416-HSP42-ΔNTD-GFP | *HSP42-Δ(1-726)-GFP-HIS3MX6*; YCp, *URA3*, *AmpR* | This study |
| pRS416-HSP42-ΔPrLD-GFP | *HSP42-Δ(1-297)-GFP-HIS3MX6*; YCp, *URA3*, *AmpR* | This study |
| pRS416-HSP42-ΔIDD-GFP | *HSP42-Δ(298-726)-GFP-HIS3MX6*; YCp, *URA3*, *AmpR* | This study |
| pRS416-HSP42-ΔACD-GFP | *HSP42-Δ(727-1041)-GFP-HIS3MX6*; YCp, *URA3*, *AmpR* | This study |
| pRS416-HSP42-ΔCTD-GFP | *HSP42-Δ(1042-1125)-GFP-HIS3MX6*; YCp, *URA3*, *AmpR* | This study |
| pRS416-HSP42-GFP(C48S,C70S) | *HSP42-GFP(142–144, 208–210:TGC → TCT)-HIS3MX6*; YCp, *URA3*, *AmpR* | This study |
| pRS416-HSP42-C127S-GFP(C48S,C70S) | *HSP42(379–381:TGT → TCG)-GFP(142–144, 208–210:TGC → TCT)-HIS3MX6*; YCp, *URA3*, *AmpR* | This study |
| pRS416-HSP42-ΔNTD-GFP(C48S,C70S) | *HSP42-Δ(1-726)-GFP(142–144, 208–210:TGC → TCT)-HIS3MX6*; YCp, *URA3*, *AmpR* | This study |
| pRS416-HSP42-ΔPrLD-GFP(C48S,C70S) | *HSP42-Δ(1-297)-GFP(142–144, 208–210:TGC → TCT)-HIS3MX6*; YCp, *URA3*, *AmpR* | This study |
| pRS416-HSP42-ΔPrLD-C127S-GFP(C48S,C70S) | *HSP42-Δ(1-297)-HSP42(379–381:TGT → TCG)-GFP(142–144, 208–210:TGC → TCT)-HIS3MX6*; YCp, *URA3*, *AmpR* | This study |
| pRS416-HSP42-ΔIDD-GFP(C48S,C70S) | *HSP42-Δ(298-726)-GFP(142–144, 208–210:TGC → TCT)-HIS3MX6*; YCp, *URA3*, *AmpR* | This study |

| | | |
|---|---|---|
| pRS416-HSP42-ΔACD-GFP(C48S,C70S) | *HSP42-Δ(727-1041)-GFP(142–144, 208–210:TGC → TCT)-HIS3MX6*; YCp, *URA3*, *AmpR* | This study |
| pRS416-HSP42-ΔCTD-GFP(C48S,C70S) | *HSP42-Δ(1042-1125)-GFP(142–144, 208–210:TGC → TCT)-HIS3MX6*; YCp, *URA3*, *AmpR* | This study |
| pIAU-HSP42prom-GFP | *HSP42prom(500)-GFP-ADH1ter*, YIp, *URA3*, *AmpR* | This study |
| pET28a-HSP42-6xHis | *HSP42-6xHis; KanR* | This study |
| pET28a-HSP42-C127S-6xHis | *HSP42(379–381:TGT → TCG)-6xHis; KanR* | This study |
| pET28a-HSP42-ΔPrLD-6xHis | *HSP42-Δ(1-297)-6xHis; KanR* | This study |
